# Condensin controls cellular RNA levels through the accurate segregation of chromosomes instead of directly regulating transcription

**DOI:** 10.1101/330654

**Authors:** Clémence Hocquet, Xavier Robellet, Laurent Modolo, Xi-Ming Sun, Claire Burny, Sara Cuylen-Haering, Esther Toselli, Sandra Clauder-Münster, Lars M. Steinmetz, Christian H. Haering, Samuel Marguerat, Pascal Bernard

**Affiliations:** CNRS Laboratory of Biology and Modelling of the Cell (LBMC), 46 Allée d’Italie, 69007, Lyon, France; Université de Lyon, ENSL, UCBL, 46 Allée d’Italie, 69007, Lyon, France; MRC London Institute of Medical Sciences (LMS), Du Cane Road, London W12 0NN, UK; Institute of Clinical Sciences (ICS), Faculty of Medicine, Imperial College London, Du Cane Road, London W12 0NN, UK; Cell Biology and Biophysics Unit, Structural and Computational Biology Unit, European Molecular Biology Laboratory, Heidelberg, Germany; Genome Biology Unit, European Molecular Biology Laboratory, Heidelberg, Germany

## Abstract

Condensins are genome organisers that shape chromosomes and promote their accurate transmission. Several studies have also implicated condensins in gene expression, although the mechanisms have remained enigmatic. Here, we report on the role of condensin in gene expression in fission and budding yeasts. In contrast to previous studies, we provide compelling evidence that condensin plays no direct role in the maintenance of the transcriptome, neither during interphase nor during mitosis. We further show that the changes in gene expression in post-mitotic fission yeast cells that result from condensin inactivation are largely a consequence of chromosome missegregation during anaphase, which notably depletes the RNA-exosome from daughter cells. Crucially, preventing karyotype abnormalities in daughter cells restores a normal transcriptome despite condensin inactivation. Thus, chromosome instability, rather than a direct role of condensin in the transcription process, changes gene expression. This knowledge challenges the concept of gene regulation by canonical condensin complexes.

## Introduction

Structural Maintenance of Chromosomes (SMC) complexes are ring-shaped ATPases, conserved from bacteria to human, which shape chromosomes and ensure their accurate transmission during cell division (Hirano, 2016; Uhlmann, 2016). Eukaryotes possess three distinct SMC protein complexes, named condensins, cohesin and SMC5/6. Condensins structure and condense chromosomes, cohesin ensures sister-chromatid cohesion and organises topological domains in the genome during interphase, and SMC5/6 promotes proper DNA replication and repair (Hirano, 2016; Uhlmann, 2016). A large body of in vivo and in vitro studies has substantiated the idea that SMC complexes shape the genome and preserve its integrity by encircling DNA helixes and, at least partly, by extruding loops of DNA (Ganji et al., 2018; Gibcus et al., 2018; Hirano, 2016; Uhlmann, 2016). Besides organising chromosomes, cohesin and condensins have also been widely implicated in the control of gene expression, raising the idea that the two complexes link gene expression to chromosome architecture (Dowen and Young, 2014). Yet, while our understanding of gene regulation by cohesin has progressed during the last decade (Merkenschlager and Nora, 2016), the mechanisms through which condensins impact on gene expression have remained largely enigmatic.

Condensins have been best characterised as the key drivers of the assembly of mitotic chromosomes (Gibcus et al., 2018; Hirano, 2016; Robellet et al., 2017). The profound reorganisation of chromatin fibres into compact and individualised rod-shaped chromosomes, that marks the entry into mitosis, is essential for the accurate transmission of the genome during anaphase. When condensins are impaired, sister-centromeres migrate towards the opposite pole of the mitotic spindle upon anaphase onset, but chromosome arms remain entangled and hence fail to separate, leading to the formation of sustained anaphase chromatin bridges and DNA breakage in telophase and in post-mitotic cells (Cuylen et al., 2013; Hyun-Soo Kim et al., 2009; Samoshkin et al., 2012; Sutani et al., 1999; Toselli-Mollereau et al., 2016; Woodward et al., 2016).

Like all SMC complexes, condensins are made of two ATPases, called SMC2^Cut14^ and SMC4^Cut3^ (fission yeast names are indicated in superscript), associated with three non-SMC subunits that regulate the ATPase activity of the holocomplex and govern its association with DNA (Kschonsak et al., 2017). Most eukaryotes possess two condensins, called condensin I and II, which are composed of a same SMC2/SMC4 heterodimer but are associated with two different sets of regulatory subunits (Hirano, 2016; Robellet et al., 2017). Despite their structural and functional similarities, the dynamics of association of condensin I and II with chromosomes differ (Walther et al., 2018). Condensin II is nuclear during interphase and enriched on chromosomes from prophase until telophase. Condensin I, in contrast, is mostly cytoplasmic during interphase and associates with chromosomes from prometaphase until telophase. A third condensin variant called the Dosage Compensation Complex (DCC) has been described in the worm *Caenorhabditis elegans*, which associates with the two X chromosomes in hermaphrodite animals and halves the expression of X-linked genes by reducing the occupancy of RNA polymerase II (RNA Pol II) (Kruesi et al., 2013). Yeasts, in contrast, possess a unique condensin complex, similar in term of primary sequence to condensin I.

Condensin I and II have been implicated in the control of gene expression in various organisms, ranging from yeasts to human. In the budding yeast *Saccharomyces cerevisiae*, the silencing of heterochromatic mating type genes and the position effect exerted by repetitive ribosomal DNA are both alleviated when condensin is impaired (Bhalla et al., 2002; Wang et al., 2016). Likewise, in the fission yeast *Schizosaccharomyces pombe*, the SMC4^Cut3^ subunit of condensin is needed to repress tRNAs genes as well as reporter genes inserted into the pericentric DNA repeats that are coated with heterochomatin (He et al., 2016; Iwasaki et al., 2010). It remains unclear, however, in which cell cycle phase and through which mechanism budding and fission yeast condensins regulate the expression of such diverse genes in such different genomic contexts. In *C. elegans*, depletion of condensin II is linked to an increase in the expression of at least 250 genes, but this effect does not correlate with the occupancy of condensin II on chromosomes (Kranz et al., 2013). In *Drosophila melanogaster*, condensin I has been implicated in the repression of homeotic genes (Lupo et al., 2001), whereas condensin II has been reported to take part in the production of antimicrobial peptides (Longworth et al., 2012). Murine peripheral T cells that express a mutant version of the condensin II CAP-H2 regulatory subunit, exhibit a decreased compaction of chromatin and an increased expression of the proliferative gene *Cis* (Rawlings et al., 2011), suggesting a possible mechanistic relationship between condensin-mediated chromosome organisation and gene expression. However, another study reported that the same CAP-H2 mutation led to only subtle effects on the transcriptome of precursor thymocytes, which might be caused by chromosomal instability (Woodward et al., 2016).

Perhaps more consistent with a direct role, condensin I was found associated with active promoters during M phase in chicken DT40 cells, and its depletion prior to mitotic entry coincided with a decreased expression during the subsequent G1 phase of a subset of genes to which it was bound to (Kim et al., 2013). Similarly, cohesin and condensin II have been detected at super-enhancers in rapidly proliferating mouse embryonic stem cells, and depleting condensin II has been associated with a reduced expression of cell-identity genes driven by these super-enhancers (Dowen et al., 2013). Yuen et al. observed a similar effect on the expression of highly-expressed housekeeping genes in mouse embryonic stem cells and human embryonic kidney cells (Yuen et al., 2017). Finally, it has been reported that not only condensin II, but also condensin I, binds enhancers activated by β-estradiol during interphase in human MCF7 breast adenocarcinoma cells, and that the depletion of condensin I or II led to a reduced transcription of oestrogen-activated genes (Li et al., 2015). Intriguingly, the same enhancers where also found occupied by cohesin and to rely upon this complex to drive gene expression (Li et al., 2013).

All these studies tend to support the idea that condensin I and II play an important and evolutionarily-conserved role in gene expression, through which they impinge on cell identity, cell proliferation and, possibly, also immunity. Yet, no conclusive evidence has been provided thus far as to how condensin I and II might achieve this function. Mitotic chromosomes conserve considerable chromatin accessibility, similar to interphase chromatin (Hihara et al., 2012), and DNA remains accessible to transcription factors even in mitotic chromosomes that have been structured by condensin complexes (Chen et al., 2005; Palozola et al., 2017). Thus, it remains unclear whether and how mechanisms related to chromosome condensation could possibly underlie condensin-mediated gene regulation. Furthermore, given that the loss or gain of chromosomes is sufficient to alter gene expression (Sheltzer et al., 2012), it is crucial to determine to which extent the role attributed to condensin I and II in the control of gene expression is mechanistically different from, or related to, the assembly and segregation of chromosomes during mitosis.

Gene expression can be controlled at the transcriptional level, by changing the activity and/or the occupancy of RNA polymerases, as exemplified by condensin^DCC^ (Kruesi et al., 2013). It can also be controlled at the co-or post-transcriptional level by modulating the half-life of transcripts (Buhler et al., 2007; Harigaya et al., 2006). The RNA-exosome is a conserved ribonuclease complex that ensures the maturation and the controlled degradation of a plethora of RNAs in the cell, including, for example, defective RNAs or cryptic unstable non-coding RNAs (Kilchert et al., 2016). The RNA-exosome consists of nine core subunits associated with the RNase Dis3 in the cytoplasm plus a second RNase, called Rrp6, in the nucleus (Kilchert et al., 2016). The mechanisms behind target recognition and processing or degradation modes selection by the RNA-exosome are not fully understood. Cofactors, such as the TRAMP poly(A)polymerase complex, stimulate the RNase activity of the RNA-exosome and specify its targets (Kilchert et al., 2016). Rrp6 has been found associated with chromosomes at actively transcribed genes (Andrulis et al., 2002), leading to the idea that the RNA-exosome can handle nuclear RNA in a co-transcriptional manner. By processing and eliminating cellular transcripts, the RNA-exosome plays a central role in proper gene expression (Kilchert et al., 2016).

To gain insights into how canonical condensins regulate genes, we investigated the role of condensin complexes in gene expression during the cell cycle, in fission and budding yeasts. In contrast with previous studies, we present here converging evidence that condensin plays no major direct role in the control of gene expression, neither in *S. pombe* nor in *S. cerevisiae*. We show that lack of condensin is associated with increased levels of mRNAs and non-coding RNAs in post-mitotic fission yeast cells, reminiscent of other organisms, but that this effect is indirect: RNAs accumulate in condensin mutant cells because the missegregation of the rDNA during anaphase dampens RNA degradation by the nucleolar RNA-exosome in post-mitotic cells. The discovery that budding and fission yeast condensins contribute to proper gene expression by maintaining chromosome stability during cell divisions, and not through a direct impact on gene transcription, challenges the widespread idea that condensin I and condensin II are direct regulators of gene expression.

## Results

### RNA-exosome-sensitive transcripts accumulate when condensin is impaired in fission yeast

To assess the role of condensin in gene expression in fission yeast, we compared the transcriptomes of wild-type and mutant cells in which the SMC2^Cut14^ ATPase subunit of condensin was inactivated by the thermosensitive *cut14-208* mutation (Saka et al., 1994). Cells growing asynchronously at permissive temperature were shifted to the restrictive temperature of 36°C for 2.5 hours (one cell doubling) to inactivate SMC2^Cut14^ and their transcriptomes determined using strand-specific RNA-seq. We identified 306 transcripts that were differentially expressed by log2 fold changes superior to 0.5 or inferior to −0.5 (p value ≤ 0.05) in the *cut14-208* condensin mutant compared to wild type (Fig. 1A). The vast majority (98.5%; n=302/306) exhibited an increased steady-state level in the mutant. We confirmed the increase for six example RNAs by RT-qPCR, using either *act1 or nda2* mRNA levels as internal controls (Fig. 1B and Fig. S1A). Thus, gene expression is altered, and mainly increased, in dividing condensin mutant *cut14-208* cells.

**Figure 1.**
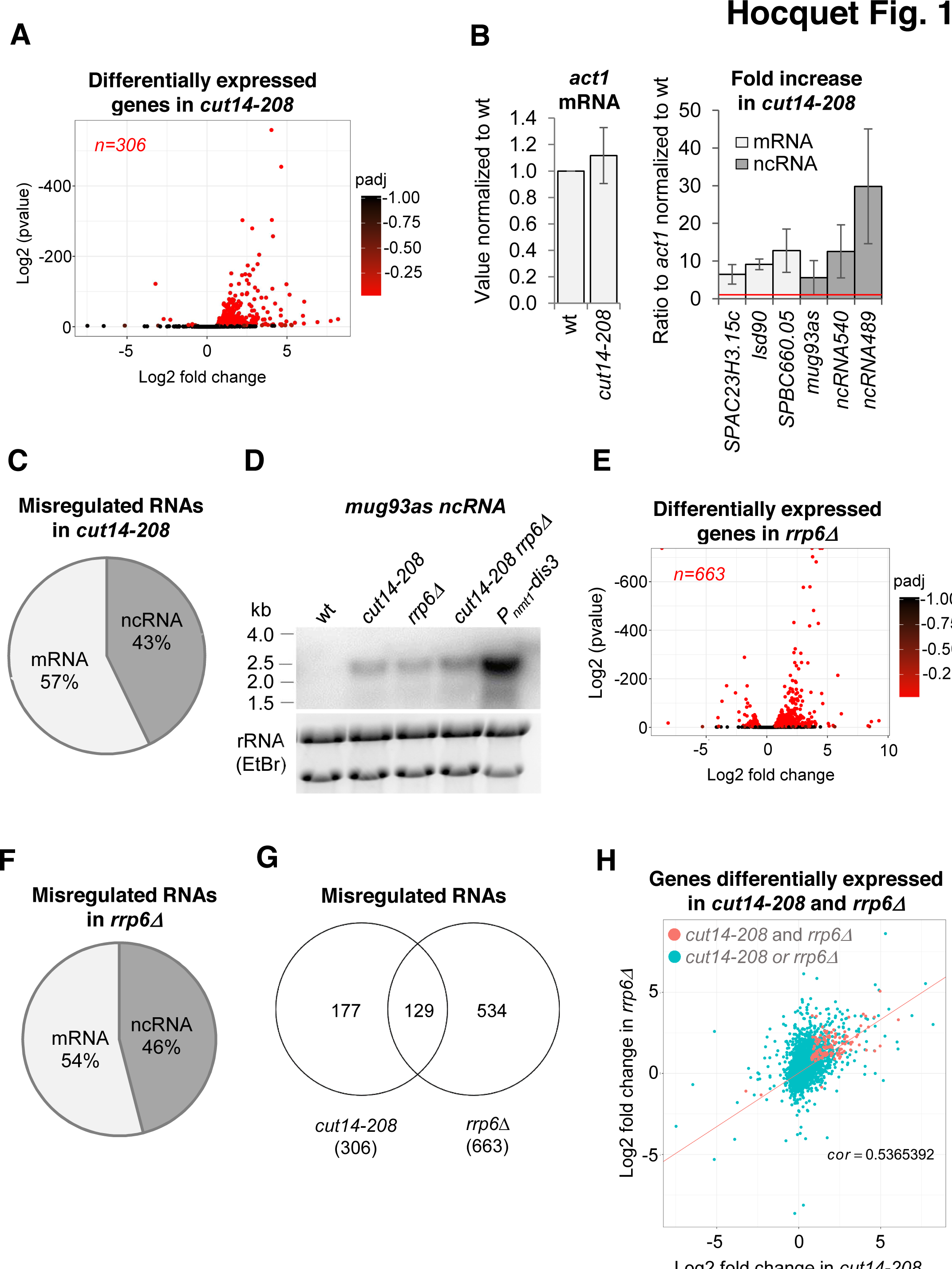
The condensin loss-of-function mutant *cut14-208* accumulates RNA-exosome-sensitive transcripts. **A.** Volcano plot of RNA levels measured by strand-specific RNA-seq in the *cut14-208* condensin mutant after 1 cell doubling at 36°C, from biological triplicates. Genes exhibiting a log2 fold change superior to 0.5 or inferior to −0.5 with an adjusted P-value (padj) <= 0.05 are indicated in red. **B.** RT-qPCR validation. Total RNA from cells grown at 36°C for 2.5 hours was reverse-transcribed in the presence or absence of Reverse Transcriptase (RT) and cDNAs were quantified by qPCR. Shown are the averages and standard deviations (SDs) measured from 3 biological replicates. **C.** Misregulated RNA in *cut14-208*. **D.** Northern blot analysis of the non-coding RNA *mug93as*. Cells were shifted at 36°C for 1 cell doubling and total RNA probed for *mug93as* level. Ribosomal RNA (rRNA) stained with ethidium bromide (EtBr) was used as loading control. **E.** Volcano plot of RNA levels measured by RNA-seq from biological triplicates of the *rrp6Δ* mutant after 1 cell doubling at 36°C. Genes exhibiting a log2 fold change superior to 0.5 or inferior to −0.5 with an adjusted P-value (padj) <= 0.05 are indicated in red. **F.** Misregulated RNA in *rrp6Δ*. **G.** Genes misregulated in *cut14-208* and *rrp6Δ*. **H.** Comparison plots between the transcriptomes of *cut14-208* and *rrp6Δ*. Genes differentially expressed in both mutants are highlighted in red. The correlation coefficient has been calculated for all genes.

Of the 306 misregulated transcripts, 57% were mRNAs and the remaining 43% were non-protein coding RNAs (ncRNA) (Fig. 1C). We found histone mRNAs amongst the upregulated transcripts, confirming previous observations (Kim et al., 2013). However, the analysis of all increased mRNAs revealed no enrichment for a specific gene ontology (GO) term, which suggests that condensin does not regulate the expression a particular family of protein-coding genes. In contrast, ncRNAs, which represent 22% of the transcription units in the fission yeast genome (n = 1522/6948), were significantly enriched in the population of transcripts up-regulated in the *cut14-208* mutant (p<0.001, Chi-square test). Since most ncRNAs are maintained at a low level by the nuclear RNA-exosome (Wilhelm et al., 2008), their controlled degradation might be compromised in the mutant. We tested this hypothesis by Northern blotting using the antisense ncRNA *mug93-antisense-1* (*mug93as*) as a representative example. As shown in Figure 1D, *mug93as* was barely detectable in a wild-type background but accumulated in cells lacking Rrp6 or defective for Dis3, as expected if it were degraded by the RNA-exosome. Remarkably, *mug93as* also accumulated in *cut14-208* mutant cells, reaching levels reminiscent of the *rrp6Δ* control. Furthermore, chromatin immunoprecipitation (ChIP) against RNA Pol II revealed no change in RNA Pol II occupancy at neither the *mug93as* gene, nor two additional example ncRNA genes (*ncRNA.489* and *ncRNA.540*), in *cut14-208* cells (Fig. S1B), although their ncRNA levels were increased between 5 and 30-fold (Fig. 1B and S1A). These RNAs might therefore accumulate due to impaired degradation rather than increased transcription. Taken together, these results indicate that unstable RNA species accumulate when condensin is defective.

To clearly delineate the number of transcripts targeted by the RNA-exosome that accumulate in the *cut14-208* condensin mutant, we compared the transcriptomes of *cut14-208* and *rrp6Δ* cells that had been grown in parallel and processed simultaneously for strand-specific RNA-seq analyses. We identified 663 RNAs that were differentially expressed by log2 fold changes superior to 0.5 or inferior to −0.5 (p value ≤ 0.05) in the *rrp6Δ* mutant (Fig. 1E). Of these differentially expressed RNAs, 78% were increased in levels and 22% were decreased. The population of Rrp6-sensitive RNAs was considerably enriched in ncRNAs (p<0.001, Chi-square test) (Fig. 1F). Pairwise comparison with *cut14-208* revealed that 41% of the RNAs that accumulated in the condensin mutant also accumulated in cells lacking Rrp6 (Fig. 1G-H), with no clear preference for ncRNA and mRNA transcripts (50% each). A hypergeometric test confirmed that this overlap was statistically highly significant (p = 4.6e^−55^). These data indicate that a large fraction of the transcripts that accumulate when condensin is impaired are normally targeted by the ribonuclease Rrp6.

### Read-through transcripts accumulate upon condensin inactivation

Visual inspection of the RNA-seq profile of the *cut14-208* mutant revealed a widespread increase of reads downstream of the 3’ ends of genes, suggesting defects in the termination of transcription (for an example, see the *hsp9* gene in Fig. 2A). Read-through transcripts are abnormal RNAs that are extended at their 3’ ends when RNA polymerases transcribe over Transcription Termination Sites (TTS) into downstream DNA sequences. Dis3 and Rrp6 have been reported to prevent the accumulation of read-through transcripts, although through possibly different mechanisms (Lemay et al., 2014; Zofall et al., 2009). Lemay et al. have shown that 3’-extended *hsp9* RNA accumulate in fission yeast cells when Dis3 is impaired, and to a lesser extend when Rrp6 is lacking (Lemay et al., 2014). By using RNase-H digestion of the 5’ end of *hps9* mRNA and northern blotting, we confirmed the accumulation of 3’-extended *hsp9* RNA in *dis3* and *rrp6* mutant cells (Fig. 2B). Importantly, *cut14-208* mutant cells accumulated 3’-extended *hsp9* RNA of the same length as cells depleted of Dis3 or Rrp6. Moreover, other condensin mutants, such as *cut3-477* and *cut14-180*, also accumulated read-through *hsp9* RNA (Fig. S2A), which demonstrates that the accumulation of uncleaved, 3’-extended *hsp9* transcripts is a feature of condensin inactivation.

**Figure 2.**
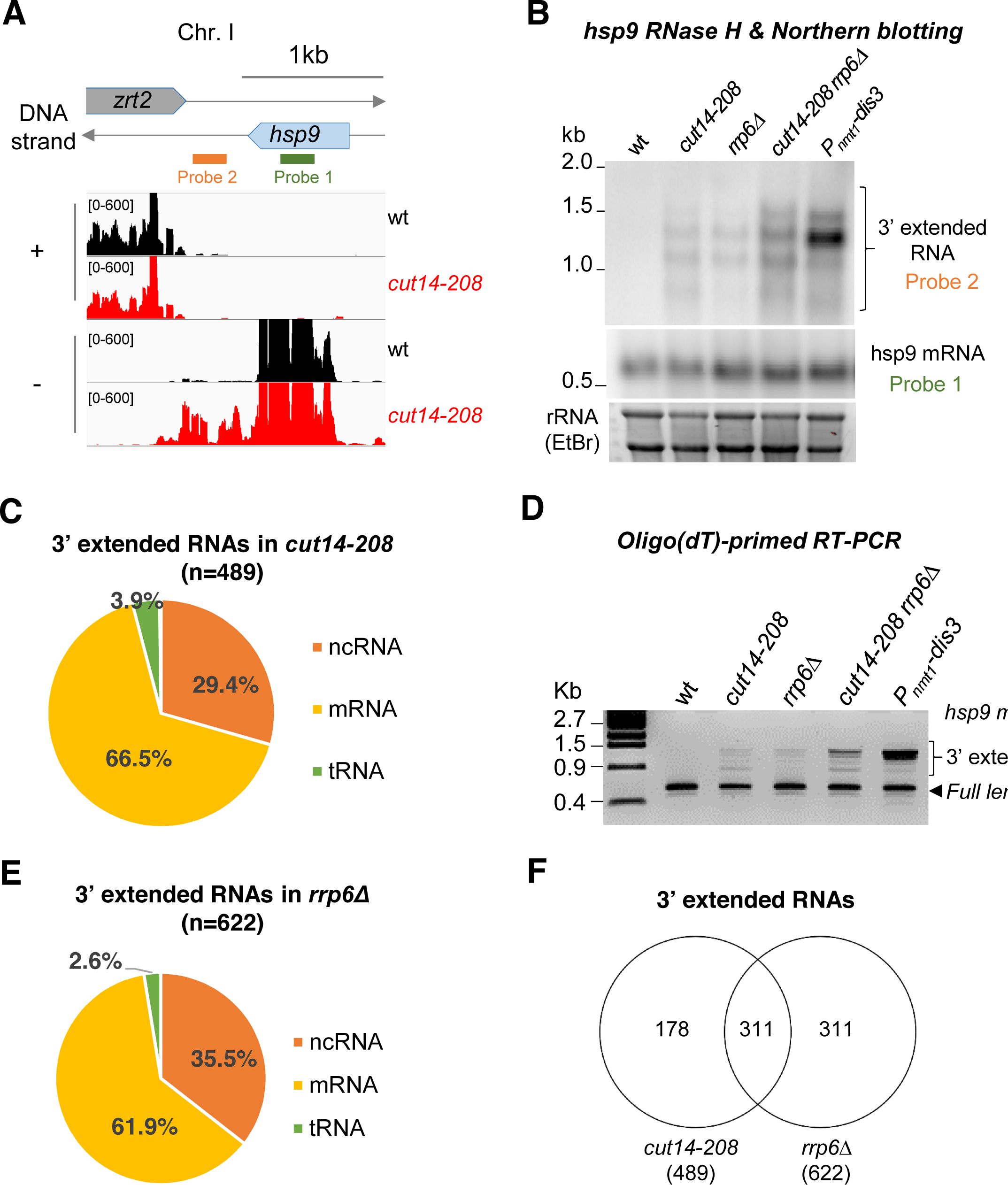
The condensin loss-of-function mutant *cut14-208* accumulates 3’-extended read-through transcripts. **A.** 3’-extended *hsp9* read-through RNA detected by strand-specific RNA-seq in *cut14-208*. **B.** Read-through *hsp9* RNA detected by RNase H digestion and Northern blotting. Total RNA from indicated strains grown at 36°C was digested by RNase H in the presence of a DNA oligonucleotide complementary to the 5’end of *hsp9* mRNA. Cleaved products were revealed by a probe hybridizing downstream the transcription termination site of *hsp9* (see probe2 in A), or within the coding sequence (probe 1, shown in A). rDNA stained with EtBr served as loading control. **C.** 3’-extended read-through RNAs in *cut14-208*. **D.** Polyadenylated RNAs detected by oligo(dT)-primed RT-PCR. Cells were grown at 36°C for 2.5 hours in PMG supplemented with 60 µM thiamine to repress *nmt1-dis3*. Total RNA was reverse transcribed using oligo(dT) primers in the presence or absence of RT. cDNA were amplified by 25 cycles of PCR using oligo(dT) and gene specific primers. Minus RT reactions produced no signal. **E.** 3’-extended read-through RNAs in *rrp6Δ*. **F.** Overlap between the sets of read-through RNAs in *cut14-208* and *rrp6Δ*.

To determine the prevalence of read-through RNAs in *cut14-208* mutant cells, we systematically searched and quantified stretches of consecutive RNA-seq reads that mapped immediately downstream the TTS of annotated genes on the same DNA strand and did not overlap with downstream genes (see Material and Methods). Using these criteria, we identified 489 transcripts, mostly mRNAs and ncRNAs, which were extended at their 3’ ends (Fig. 2C). Oligo(dT)-primed RT-PCR showed that read-through RNAs were polyadenylated (Fig. 2D and S2B), suggesting that non-canonical polyadenylation sites were more frequently used in the *cut14-208* mutant than in wild-type cells, as already described for cells lacking Dis3 (Lemay et al., 2014). GO term analysis revealed no specific feature defining the population of read-through transcripts that accumulate in *cut14-208* cells. Furthermore, only 111 of the 489 read-through transcripts were also up-regulated (Fig. S2C), which suggests that the 3’-extended RNAs were not the by-product of increased transcription.

To further investigate the molecular origin of read-through RNAs in condensin mutant cells, we compared *cut14-208* with *rrp6Δ* cells. We detected 622 read-through RNAs, again mostly mRNAs and ncRNAs, in the transcriptome of cells lacking Rrp6 (Fig. 2E), which confirms previous reports (Lemay et al., 2014; Zofall et al., 2009). Importantly, 50% of these RNAs were also extended at their 3’ends in the *cut14-208* mutant (Fig. 2F). This reinforces the idea that the function of Rrp6 might be affected in a *cut14-208* background. Note that the steady state level of the Rrp6 protein remained unchanged in *cut14-208* mutant cells (Fig. S2D). Likewise, we observed no change in the mRNA levels of RNA-exosome or TRAMP components in the condensin mutant by RNA-seq (Table S1). Collectively, these data indicate that 3’-extended read-through transcripts accumulate upon condensin inactivation in fission yeast and that this accumulation might stem from defects in RNA processing of these transcripts by Rrp6.

### Condensin is dispensable for gene expression during interphase and metaphase in fission and budding yeasts

Since condensin is regulated over the course of the cell cycle, we sought to determine the phase(s) during which condensin function is required for proper gene expression. We synchronised *cut14-208* mutant cells in early S phase at the permissive temperature, raised the temperature to 36°C to inactivate condensin and at the same time released them into the cell cycle. We then measured gene expression by RT-qPCR as cells progressed from early S phase into the cell cycle (Fig. 3A). To ensure that cells went only through a single cell cycle at 36°C, we re-arrested cells in late G1 phase by the thermosensitive mutation *cdc10-129*. Previous work had shown that 10 min at 36°C are sufficient to inactivate condensin in *cut14-208* mutant cells (Nakazawa et al., 2011). FACScan analysis of DNA content and cytological observations revealed that *cdc10-129* single mutant and *cdc10-129 cut14-208* double mutant cells completed S phase (t = 30 min after release) and progressed through G2 phase (t = 60 min) and mitosis (t = 90 min) with similar kinetics (Fig. 3B). Chromosome segregation was impaired in *the cut14-208* mutant background, as revealed by the appearance of anaphase chromatin bridges (Fig. 3B, green line), which were subsequently severed by the septum upon mitotic exit, producing the CUT phenotype (Cells Untimely Torn) (Fig. 3B, red line). Accordingly, FACScan analysis revealed the appearance of aberrant karyotypes in post-mitotic cells (t = 120 and 180 min; Fig. 3B).

**Figure 3.**
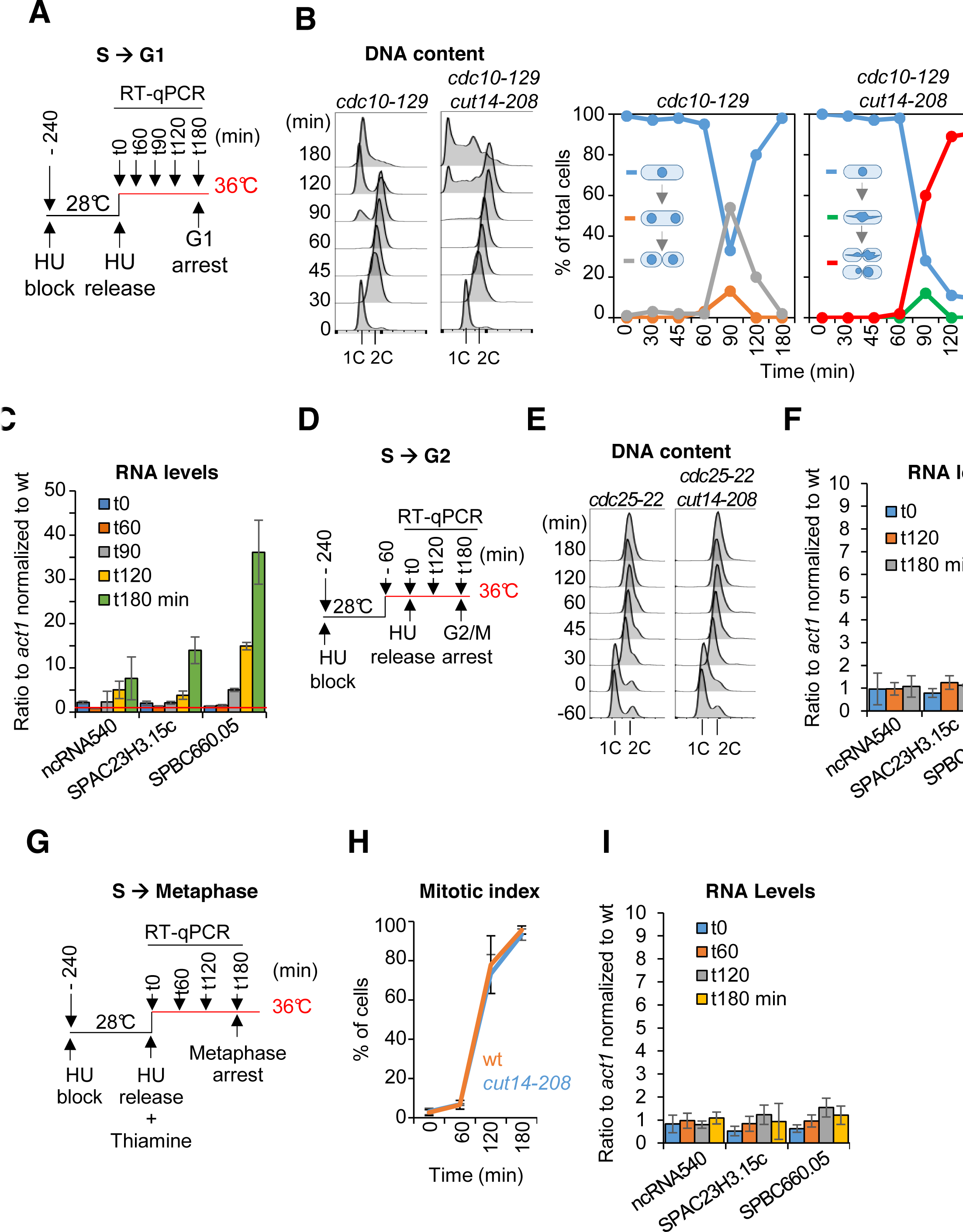
The function of condensin is dispensable for gene regulation during S and G2 phases of the cell cycle in fission yeast. **A-C.** Gene expression was assessed in synchronized *cut14-208* cells progressing from early S phase to G1 phase at the restrictive temperature**. A**. Scheme of the experiment**. B.** Left panel: FACScan analyses. Right panels: chromosome segregation and cytokinesis assessed by staining DNA with DAPI and the septum with calcofluor (n >= 100). **C.** Total RNA extracted from *cdc10-129* and *cdc10-129 cut14-208* cells shown in **B**, was reverse-transcribed in the presence or absence of RT and cDNA quantified by qPCR. Red line = 1. Shown are averages ± SDs measured from biological triplicates. **D-F.** Gene expression was assessed in synchronized *cut14-208* cells progressing from early S phase to late G2 phase at the restrictive temperature. **D.** Scheme of the experiment**. E.** FACScan analyses. **F.** Total RNA extracted from *cdc25-22* and *cdc25-22 cut14-208* cells shown in **E**, was reverse-transcribed in the presence or absence of RT and cDNA quantified by qPCR. Shown are averages ± SDs measured from biological triplicates. **G-I.** Gene expression was assessed in synchronized *cut14-208* cells progressing from early S phase to metaphase at the restrictive temperature. **G.** Scheme of the experiment. Thiamine repressed the *nmt41-slp1* gene in order to arrest cells in metaphase. **H.** Percentages of mononucleate, mitotic cells from n = 3 experiments. **I.** Total RNA extracted from *nmt41-slp1 and nmt41-slp1 cut14-208* cells shown in **H** was reverse-transcribed in the presence or absence of RT and cDNA quantified by qPCR. Shown are averages ± SDs measured from biological triplicates.

Remarkably, we measured no up-regulation of any of the three reporter RNAs that we had selected from the pool of upregulated transcripts in *cut14-208* mutants (Fig. 1B) during G2 phase in synchronized *cdc10-129 cut14-208* cells (t = 60 min, Fig. 3C). These RNAs started to accumulate, however, at t = 90 min and further increased at t= 120 min and 180 min (Fig. 3C), coincidently with the accumulation of aneuploid post-mitotic cells. This result suggests a possible requirement for condensin for proper gene expression during late mitosis or early G1 phase. To validate these results, we repeated the RT-qPCR measurements by synchronously releasing *cut14-208* cells from the early S phase block only 1 hour after shifting the temperature to 36°C and re-arrested them already at the G2/M transition using a *cdc25-22* mutation (Fig. 3D-E). In this time course experiment, we observed no up-regulation of the reporter RNAs at any time point (Fig. 3F). Similarly, RNA levels remained unchanged after a sustained arrest in metaphase at 36°C by depletion of the anaphase promoting complex (APC/C) activator Slp1 (Fig. 3G-I). We conclude that, in fission yeast, the function of condensin is dispensable for gene regulation during S and G2 phases of the cell cycle, consistent with the finding that condensin is largely displaced from the nucleus during this time (Sutani et al., 1999).

Condensin in the budding yeast *S. cerevisiae* is, in contrast, bound to chromosomes throughout the cell cycle (D’Ambrosio et al., 2008), much like the condensin II complex in metazoan cells. This raises the possibility that condensin controls interphase gene expression in this species. We first confirmed that condensin localizes to chromosomes of cells released synchronously into the cell cycle from a G1 mating pheromone arrest (Fig. S3A). Chromosome spreading and ChIP showed that condensin bound to chromosomes already during G1 phase and that its levels on chromosomes increased as cells passed through S and G2 phases (Fig. S3B-C). To inactivate condensin in budding yeast, we proteolytically cleaved the kleisin (Brn1) subunit of the condensin ring by inducing expression of a site-specific protease from tobacco etch virus (TEV) using a galactose-inducible promoter, which efficiently released condensin from chromosomes (Cuylen et al., 2011), even during G1 phase (Fig. S3D). We then compared the transcriptome of G1 phase-arrested cells after condensin cleavage to cells with intact condensin (Fig. 4A). Remarkably, solely 26 transcripts were differentially expressed by at least two-fold (Fig. 4B). To rule out that this minor effect on gene expression was the result of the G1 phase arrest, we repeated the experiment, but this time released cells after Brn1 TEV cleavage from the G1 phase arrest and re-arrested them in the subsequent M phase by addition of the spindle poison nocodazole before preparing RNA for transcriptome analysis (Fig. 4C). In this experiment, only six genes showed an up-or downregulation of two-fold or more (Fig. 4D). We conclude that condensin inactivation by releasing the complex from chromosomes has no major effects on the gene expression program of budding yeast cells.

**Figure 4.**
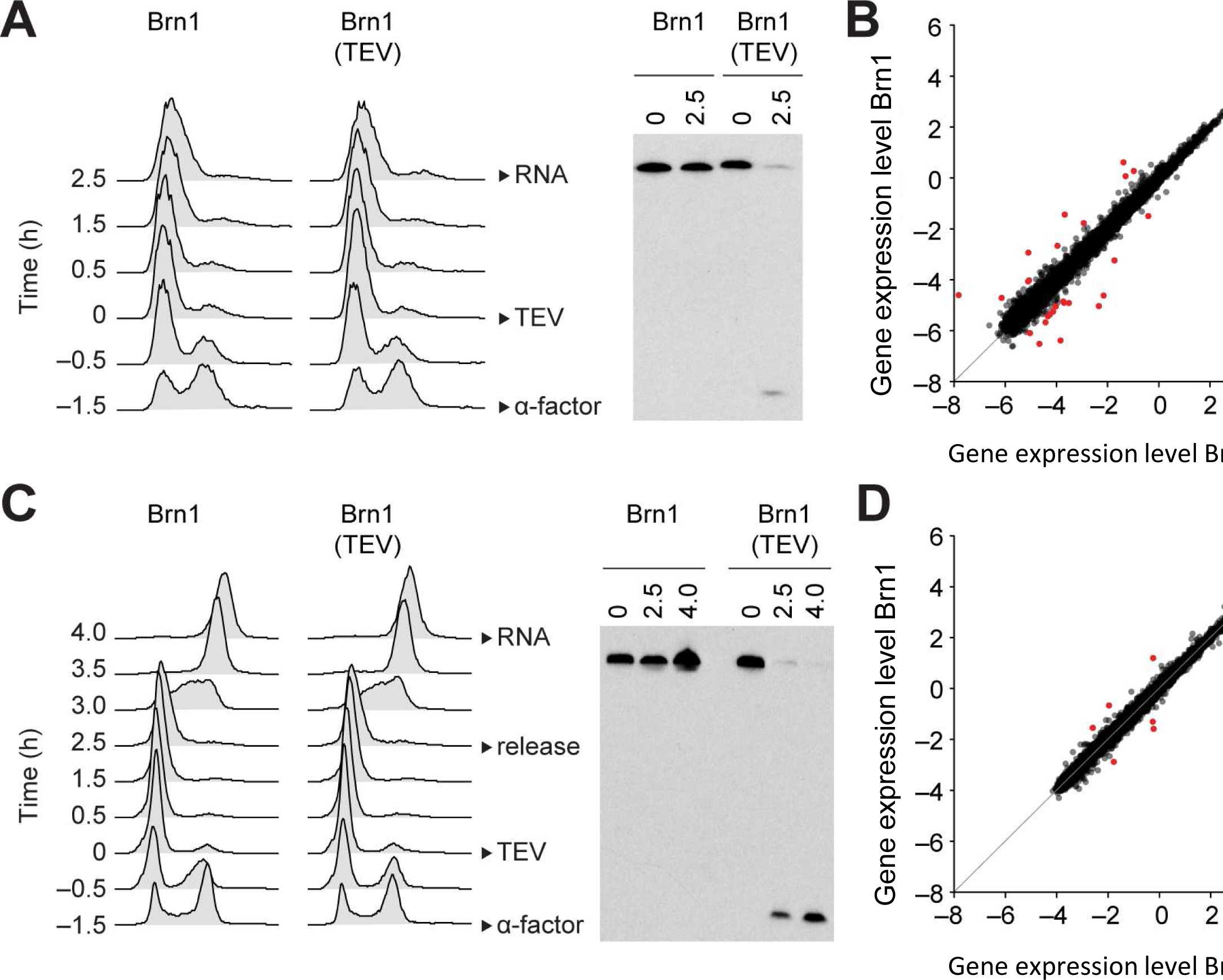
Condensin release from chromosomes has no major effects on G1 or M phase gene expression programs in budding yeast. **A.** TEV protease expression was induced in cells synchronized in G1 phase by α-factor (strains C3138 and C3139). 2.5 h after TEV induction, RNA was extracted, cDNA synthesized, labelled and hybridized to tiling arrays. Cell cycle synchronization was scored by FACScan analysis of cellular DNA content and Brn1 cleavage was monitored by western blotting against the C-terminal HA_6_ tag. **B.** Scatter plot of gene expression values of cells from **A** with cleaved or intact Brn1 (mean values of n=3 biological replicates). Red color highlights two-fold or more up-or downregulated transcripts. **C.** TEV protease expression was induced in cells synchronized in G1 phase by α-factor (strains C2335 and C2455). Cells were release into nocodazole 2.5 h after TEV induction and RNA was extracted 1.5 h later, cDNA synthesized, labelled and hybridized on tiling arrays. Cell cycle synchronization and Brn1 cleavage was monitored as in A. **D.** Scatter plot of gene expression values of cells from **C** with cleaved or intact Brn1 (mean valued of n=2 biological replicates). Red color highlights two-fold or more up-or downregulated transcripts.

### Gene expression changes in fission yeast condensin mutants are the result of a loss of genome integrity

The fact that we observed changes in transcription levels in asynchronously dividing *cut14-208* fission yeast mutant cells, but not in fission or budding yeast condensin mutants that are prevented from undergoing anaphase, raises the possibility that defects in chromosome segregation caused by condensin inactivation might be causally responsible for condensin-dependent gene deregulation. Indeed, when we arrested fission yeast *cut14-208* cells at the G2/M transition using the analogue-sensitive Cdc2asM17 kinase (Aoi et al., 2014), shifted the temperature to 36°C and then released them from the arrest to complete mitosis, we measured an increase in transcript levels after mitotic exit and the severing of chromosomes (Fig. S4).

If RNA misregulation were indeed caused by the severing of missegregated chromosomes by the cytokinetic ring, then three key hypotheses should prove correct: (1) any mutation that results in chromosome missegregation and cutting upon mitotic exit should result in an increase in levels of the same or a similar set of RNAs as the *cut14-208* mutation, (2) the amplitude of the increase in RNA levels should correlate with the prevalence of missegregation, and (3) preventing chromosome cutting in *cut14-208* cells should attenuate the changes in RNA levels. As shown in Fig. 5A, mutation of separase (*ptr4-1*) or of a subunit of the APC/C (*cut9-665*) results in chromosome missegregation and severing by the cytokinetic ring upon mitotic exit and an increase in RNA levels in a manner similar to *cut14-208*. Moreover, the amplitude of the increase in RNA levels correlated with the frequency of chromosome cutting in the different mutants (Fig. 5A). The analysis of five additional condensin mutations of increasing prevalence further confirmed this correlation (Fig. S5A-B). Furthermore, we found that RNA levels remained comparable to wild-type in *cut14-208* cells that were prevented from undergoing cytokinesis. The thermosensitive *cdc15-140* mutation prevents cytokinesis at the restrictive temperature (Balasubramanian et al., 1998). Double mutant cells *cdc15-140 cut14-208* exhibited chromatin bridges during anaphase at 36°C (Fig. 5B), indicating that *cdc15-140* did not supress the chromosome segregation defect caused by the *cut14-208* mutation. However, in the absence of a cytokinetic ring, these chromatin bridges were no longer severed upon mitotic exit, which suppressed the production of karyotypically-aberrant post-mitotic cells (Fig. 5B). Remarkably, parallel RNA-seq analysis revealed that ∼98% of the RNA up-regulated in the *cut14-208* single mutant were no-longer detected as differentially expressed in the double mutant *cut14-208 cdc15-140* (Fig. 5C-D). The suppressive effect of *cdc15-140* on *cut14-208* with respect to the accumulation of the anti-sense RNA *mug93as* is shown as an example in Fig. 5E. The production of read-through transcripts was similarly suppressed (Fig. S5C-D). Note that RNA levels remained increased in the *cdc15-140 rrp6Δ* mutant (Fig. S5E), ruling out a potential compensatory effect of *cdc15-140* on Rrp6 deficiency per se. Finally, we found that *cdc12-112,* another mutation that also impairs cytokinesis (Chang et al., 1997), equally restored normal gene expression in a *cut14-208* genetic background (Fig. S5F), confirming that cytokinesis was a driving force for the gene deregulation exhibited by the *cut14-208* mutant. Taken together, these data indicate that changes in gene expression exhibited by *cut14-208* condensin mutant cells are mostly, if not entirely, the consequence of cytokinesis when condensin is inactivated.

**Figure 5.**
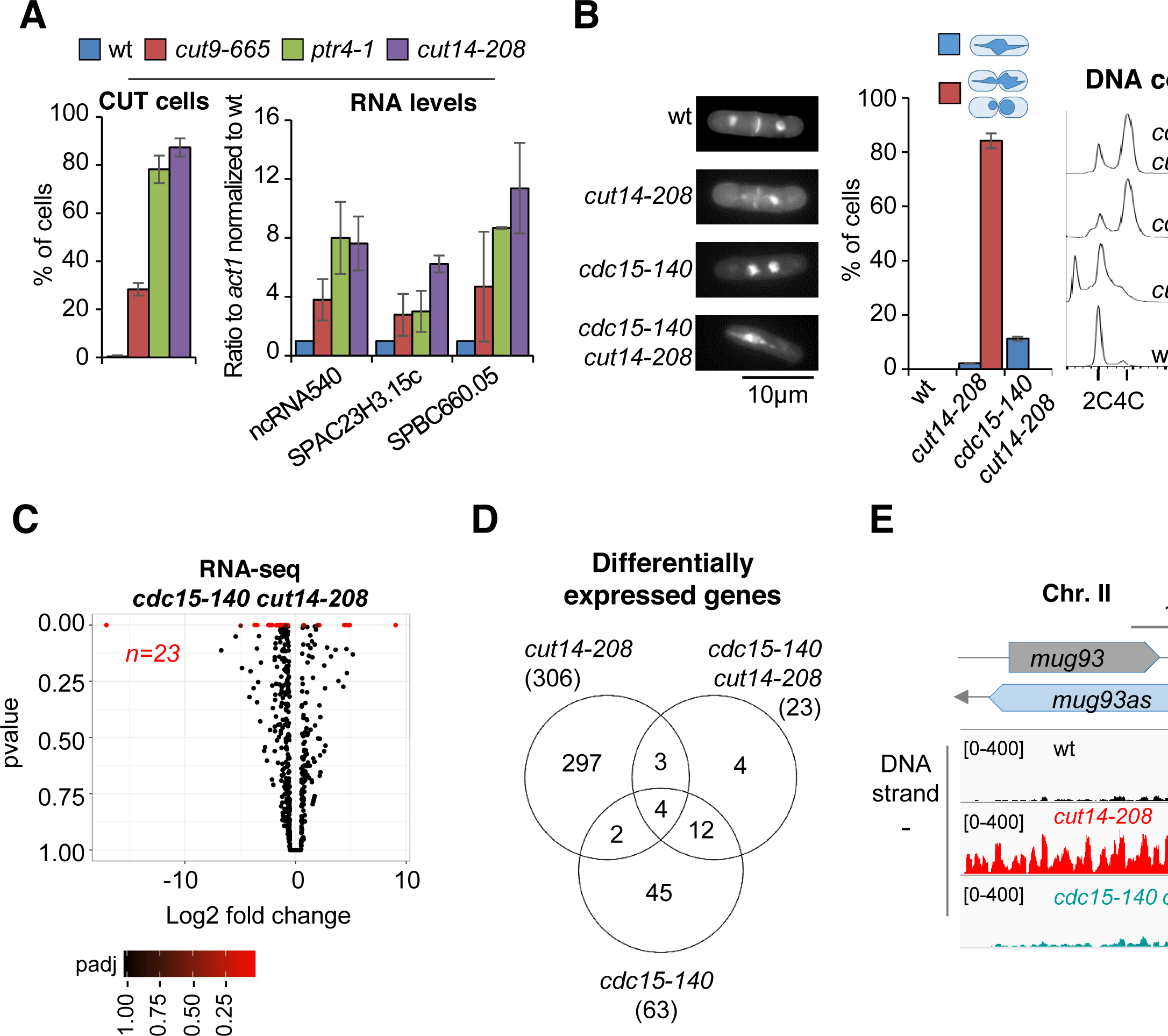
Defective mitosis underlies deregulated gene expression in the fission yeast *cut14-208* condensin mutant. **A.** Gene deregulation in mutant cells in which chromosomes are cut by the cytokinetic ring upon mitotic exit. Strains grown at 36°C for 2.5 hours were processed for cytological analysis and RT-qPCR. Right: cells were stained with DAPI and calcofluor to visualise DNA and the septum, respectively, and to quantify the frequency of chromosome cutting by the septum (CUT cells). Left: total RNA was reverse-transcribed in the presence or absence RT and cDNA quantified by qPCR. Shown are averages ± SDs calculated from 3 biological replicates. **B-E.** Preventing chromosome severing restores normal gene expression in the condensin mutant *cut14-208*. **B.** Cells were grown at 36°C for 2.5 hours and stained with DAPI and calcofluor to reveal DNA and the septum, and measure the frequency of CUT cells, or treated for FACScan analysis of DNA content. **C.** Volcano plot of RNA levels measured by strand-specific RNA-seq in the *cdc15-140 cut14-208* double mutant after 2.5 hours at 36°C, from biological triplicates. **D.** Comparative RNA-seq transcriptomic analysis from biological triplicates. **E.** RNA-seq profiles of the *mug93* ncRNA.

The severing of chromosomes by the cytokinetic ring leads to DNA damage, as revealed by the accumulation of sustained Rad22-GFP foci (Fig. S6A), but also to the formation of genomically-aberrant daughter cells, as shown by FACScan analysis of DNA contents (Fig. 3B). Both phenotypes coincide with the increase in RNA levels, and both are suppressed by *cdc15-140* (Fig. 5B and S6A). To test whether one or both might be responsible for gene deregulation in *cut14-208* cells, we analysed by RT-qPCR the impact of DNA damage upon mitotic exit, or of genomic imbalance, on gene expression in cells provided with a fully functional condensin. To damage DNA, cells were synchronized in prometaphase by using the cryosensitive mutation *nda3-KM311*, and released into mitosis in the presence of Camptothecin to induce DNA breaks during S phase, which in fission yeast overlaps with cytokinesis and septum formation. As an alternative experiment, wild-type cycling cells were treated with Zeocin. The appearance of sustained Rad22-GFP foci in cells treated with Camptothecin or with Zeocin confirmed the accumulation of DNA damage (Fig. S6B). However, RT-qPCR revealed no increase in RNA levels (Fig. S6B), arguing that DNA damage is not the main driver for gene deregulation in *cut14-208* mutant cells.

To test the impact of genomic imbalance on gene expression, we used the thermosensitive *mis6-302* mutation which disrupts kinetochore assembly, and hence accurate chromosome segregation during anaphase, leading to the production of aneuploid post-mitotic cells (Saitoh et al., 1997). Note that *mis6-302* causes neither chromatin bridges nor a CUT phenotype. FACScan analysis of *mis6-302* cells grown at the restrictive temperature confirmed the appearance of genomically-aberrant cells (Fig. 6A). Remarkably, RT-qPCR of four reporter RNAs revealed a similar increase in *mis6-302* cells as in *cut14-208* mutant cells (Fig. 6B). Therefore, this result strongly suggests that the imbalance in genomic content that results from chromosome missegregation during mitosis is a major cause of gene deregulation when condensin function is impaired.

**Figure 6.**
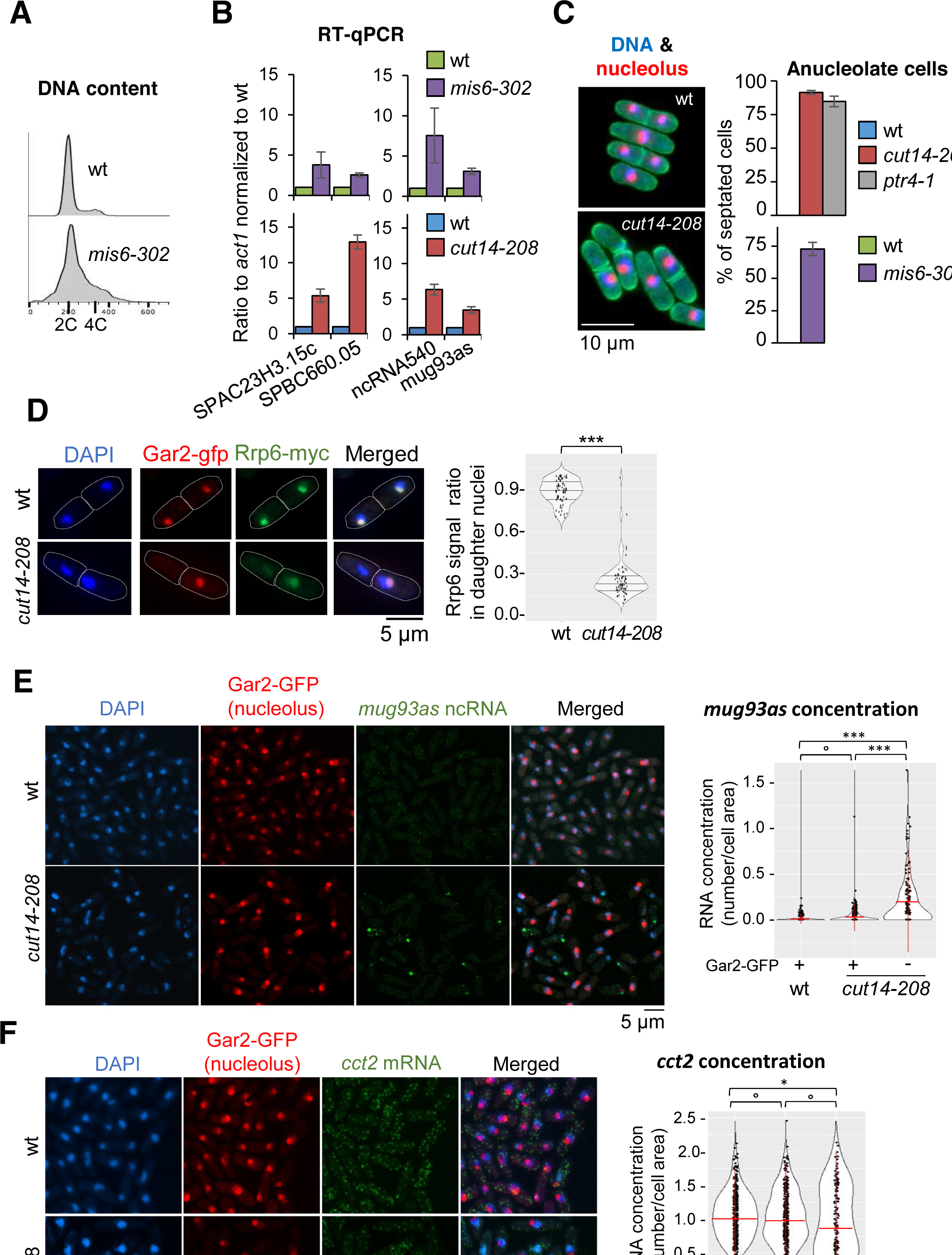
Condensin inactivation generates anucleolate daughter cells in fission yeast, which are depleted of the RNases Rrp6 and Dis3 and accumulate unstable RNA. **A-B.** The kinetochore mutation *mis6-302* and the condensin mutation *cut14-208* deregulate a same set of genes. Wildtype and *mis6-302* cells grown at 36°C for 8 hours were processed to analyse DNA content by FACScan (**A**) and RNA levels by RT-qPCR (**B**). *cut14-208* cells and the isogenic wt control grown at 36°C for 2.5 hours were used for comparison. Shown are averages ± SDs measured from biological triplicates. **C.** Non-disjunction of the rDNA in *cut14-208* and *mis6-302* cells. The nucleolar protein Gar2-mcherry was used as a marker for the rDNA and the plasma membrane protein Psy1-GFP to visualise cytokinesis. Mutant cells and their isogenic wt controls were grown at 36°C for 2.5h (*cut14-208*) or 8 hours (*mis6-302*), fixed and stained with DAPI. Segregation of the rDNA in daughter nuclei was measured upon mitotic exit. **D.** Rrp6 is enriched in the nucleolus, and depleted from anucleolate *cut14-208* mutant cells. Indicated cells were grown at 36°C, fixed and processed for immunofluorescence against Gar2-GFP and Rrp6-myc. DNA was stained with DAPI. Right panel shows the ratio of Rrp6-myc signals measured within daughter nuclei in septated cells. **E-F.** The non-coding RNA *mug93as* accumulate in anucleolate *cut14-208* cells. Cells of indicated genotype and expressing Gar2-GFP were grown at 36°C for 2.5 hours, fixed and processed for single molecule RNA FISH using probes complementary to the ncRNA *mug93as* (**E**) or the mRNA *cct2* (**F**). Box and whiskers plots show quantifications of RNA spots in *cut14-208* compared to wt, and in nucleolate compared to anucleolate mutant cells. ***p<0.01, *p<0.05 and °p>0.05.

### Non-disjunction of the rDNA during anaphase depletes the RNA-exosome from post-mitotic fission yeast cells and results in ncRNA accumulation

To investigate further the mechanism of gene deregulation when condensin is impaired, we looked in more details at chromosome segregation during anaphase in *cut14-208* mutant cells. It has been reported that condensin plays a major role in the segregation of the rDNA in late anaphase in budding yeast (D’Ambrosio et al., 2008; Freeman et al., 2000). We reached the same conclusion in fission yeast. When we scored the segregation of a GFP-labelled version of the rDNA-binding protein Gar2 in *cut14-208* mutant cells, we found that sister rDNA copies failed to separate during anaphase, frequently resulting in the formation of anucleolate daughter cells (Fig. 6C). The *ptr4-1* and *mis6-302* mutants also exhibited a high rate of anucleolate cell formation (Fig. 6C) and the *cdc15-140* mutation supressed the formation of anucleolate cells in *cut14-208* mutants (Fig. S6C). Thus, the inability to properly segregate the rDNA during anaphase correlates with the accumulation of mRNAs and ncRNAs targeted by the RNA-exosome.

Live imaging of Rrp6 and Dis3 has revealed that the two RNases are enriched in the nucleolus of fission yeast cells (Yamanaka et al., 2010). Given the overlap between the differential transcriptomes of *cut14-208* and *rrp6Δ* mutants, we hypothesised that rDNA non-disjunction might alter the localisation of Rrp6 in daughter cells upon mitotic exit. Co-immunostaining of Rrp6 tagged with a myc epitope and Gar2 tagged with GFP confirmed the nuclear localisation of Rrp6 and its marked enrichment within the nucleolus (Fig. 6D). In a wild-type background, Rrp6 appeared evenly distributed between nuclei in post-mitotic daughter cells (median signal ratio ∼1). In sharp contrast, the amount of Rrp6 was markedly reduced in *cut14-208* anucleolate daughter cells compared to their nucleolate counterparts (median signal ratio ∼0.25; Fig. 6D). We observed a similar asymmetrical distribution of Dis3 tagged with an HA epitope in post-mitotic *cut14-208* cells (Fig. S6D). Thus, condensin deficiency leads to rDNA non-disjunction during anaphase and the production of anucleolate post-mitotic cells, which are depleted of Rrp6 and Dis3.

The depletion of Rrp6 and Dis3 provided a plausible cause for the accumulation of exosome-sensitive RNA in asynchronously dividing *cut14-208* condensin mutant cells. If it were the case, exosome-sensitive RNAs should accumulate preferentially in anucleolate *cut14-208* cells. Single molecule RNA-FISH showed that the ncRNA *mug93as* produced very faint signals in wild-type cells (Fig. 6E), consistent with its active degradation by the RNA-exosome (Fig. 1D). On the contrary, *mug93as* levels were considerably higher in *cut14-208* mutant cells, which confirms our previous RNA-seq and RT-qPCR data (Fig. 6E). Crucially, *mug93as* RNA accumulated mainly in anucleolate cells devoid of Gar2-GFP, whereas a control mRNA (*cct2*) was evenly distributed in *cut14-208* mutant cells (Fig. 6F). We conclude that condensin loss-of-function in fission yeast leads to the formation of anucleolate cells that accumulate RNA-exosome-sensitive transcripts.

## Discussion

Condensin I and II have been implicated in the control of gene expression in a wide range of organisms, but it has remained unclear what aspect of the gene expression program they affect. Here, we challenge the idea that condensin complexes directly regulate transcription by providing compelling evidence that the functional integrity of condensin is dispensable for the maintenance of proper gene expression during interphase and even mitosis in fission and budding yeasts. Consistent with previous studies, we show that condensin deficiency alters the transcriptome of post-mitotic cells in fission yeast. However, we further show that this effect is mostly, if not entirely, the indirect consequence of chromosome missegregation during anaphase, which notably depletes the RNA-exosome from daughter cells. Our findings therefore indicate that condensin plays no direct role in the control of gene expression in fission and budding yeasts, but is essential for the maintenance of proper gene expression by contributing to the accurate segregation of chromosomes during mitosis.

Strand-specific RNA seq analysis revealed that *cut14-208* condensin mutant cells accumulate mRNA, ncRNA and 3’-extended read-through transcripts. Additional condensin mutants such as *cut3-477, cut3-m26, cut14-90* and *cut14-180* exhibited similar increases in RNA levels, arguing that condensin takes part in proper gene expression in fission yeast, alike in other organisms. The population of *cut14-208* condensin-mutant cells that exhibited increased RNA levels went through G2, M, G1, and S phases at the restrictive temperature in an asynchronous manner before their RNA was extracted and analysed. However, when *cut14-208* cells were synchronised during the S or G2 phases or even during metaphase, RNA levels remained unchanged. Moreover, the prevalent suppressive effect of *cdc15-140* upon *cut14-208* in term of changes in gene expression, argues (1) that condensin impinges on gene expression in a cytokinesis-dependent manner, and therefore in an indirect manner, and (2) that condensin deficiency has in itself no predominant impact on the steady state level of RNAs in post-mitotic cells. Thus, although we cannot rule out the possibility that subtle impacts of condensin on the dynamics of transcription might have escaped our detection, our data strongly indicate that condensin plays no major direct role in the maintenance of proper gene expression during interphase and even mitosis in fission yeast. Concordantly, we show that in budding yeast too, condensin is largely dispensable for the maintenance of gene expression during the G1 phase or during metaphase. Importantly, our results are in perfect agreement with recent data supporting the idea that the global transcriptional program of *S. cerevisiae* is largely insensitive to condensin depletion (Paul et al., 2017). Note that the separation of the budding and fission yeast lineages is thought to have occurred ∼ 420 million years ago, which makes them as different from each other as either is from animals (Sipiczki, 2000). Thus, our results provide compelling evidence that, in two evolutionarily distant organisms, condensin complexes play no major direct role in the maintenance of an established gene expression program.

The corollary is that the increased RNA levels exhibited by fission yeast *cut14-208* mutant cells must be the indirect consequence of a failure during late mitosis caused by a lack of condensin activity. This conclusion is in agreement with our observation that RNA levels increase coincidently with mitotic exit in synchronized condensin-mutant cells. In line with this, we provide evidence that the non-disjunction of the duplicated copies of the rDNA during anaphase constitutes a major source of gene deregulation in condensin mutant cells. (1) We show that sister-rDNA almost systematically fail to disjoin during anaphase in *cut14-208* cells, leading to the production of anucleolate daughter cells. (2) Rrp6 and Dis3 are enriched in the nucleolus in wild-type cells, but largely depleted from anucleolate *cut14-208* cells. Rrp6 accumulates predominantly on chromosomes during mitosis in Drosophila (Graham et al., 2009), suggesting that the bulk of Rrp6 and Dis3 might similarly co-segregate with the nucleolus during anaphase in fission yeast. (3) We show by RNA-seq, RT-qPCR and Northern blotting that the depletion of Rrp6 and Dis3 from anucleolate condensin mutant cells coincides with an increased steady state level of RNA-exosome-sensitive transcripts. Note that the slight additive effect exhibited by the *cut14-208* and *rrp6Δ* mutations with respect to the accumulation of *mug93as* (Fig. 1E) and of 3’-extented *hsp9* RNA (Fig. 2B) is consistent with the co-depletion of Rrp6 and Dis3 from anucleolate *cut14-208* mutant cells. (4) We further show by RNA-FISH that an RNA-exosome sensitive transcript, the antisense RNA *mug93as,* accumulates principally if not exclusively in anucleolate condensin mutant cells, as expected from an impaired degradation. (5) Importantly, supressing the production of anucleolate daughter cells by the cytokinesis mutation *cdc15-140*, in a *cut14-208* mutant background, is sufficient to restore an almost normal transcriptome, despite condensin remaining impaired. Mitosis being closed in fission yeast, when cytokinesis is prevented, anaphase chromatin bridges and the nucleolus collapse into a single nucleus upon mitotic exit. (6) Finally, we show that, reciprocally, generating anucleolate and aneuploid daughter cells by disrupting chromosome segregation by the kinetochore mutation *mis6-302*, is sufficient to trigger the accumulation of RNA-exosome-sensitive transcripts, alike *cut14-208*, but without mutating condensin.

Based on these observations, we propose the model depicted in Figure 7. Condensin deficiency impairs chromosome segregation and notably leads to chromatin bridges and the non-disjunction of the rDNA during anaphase. Rrp6 and Dis3, which are enriched in the nucleolus, and which most likely segregate with the bulk of the rDNA during anaphase, become depleted from anucleolate daughter cells produced upon mitotic exit. This strongly reduces the activity of the RNA-exosome in anucleolate daughter cells, allowing the accumulation of RNA molecules that are normally actively degraded by the RNA-exosome, such as ncRNAs and 3’-extended RNAs. This sequence of events illustrate how condensin deficiency indirectly changes gene expression in fission yeast.

**Figure 7.**
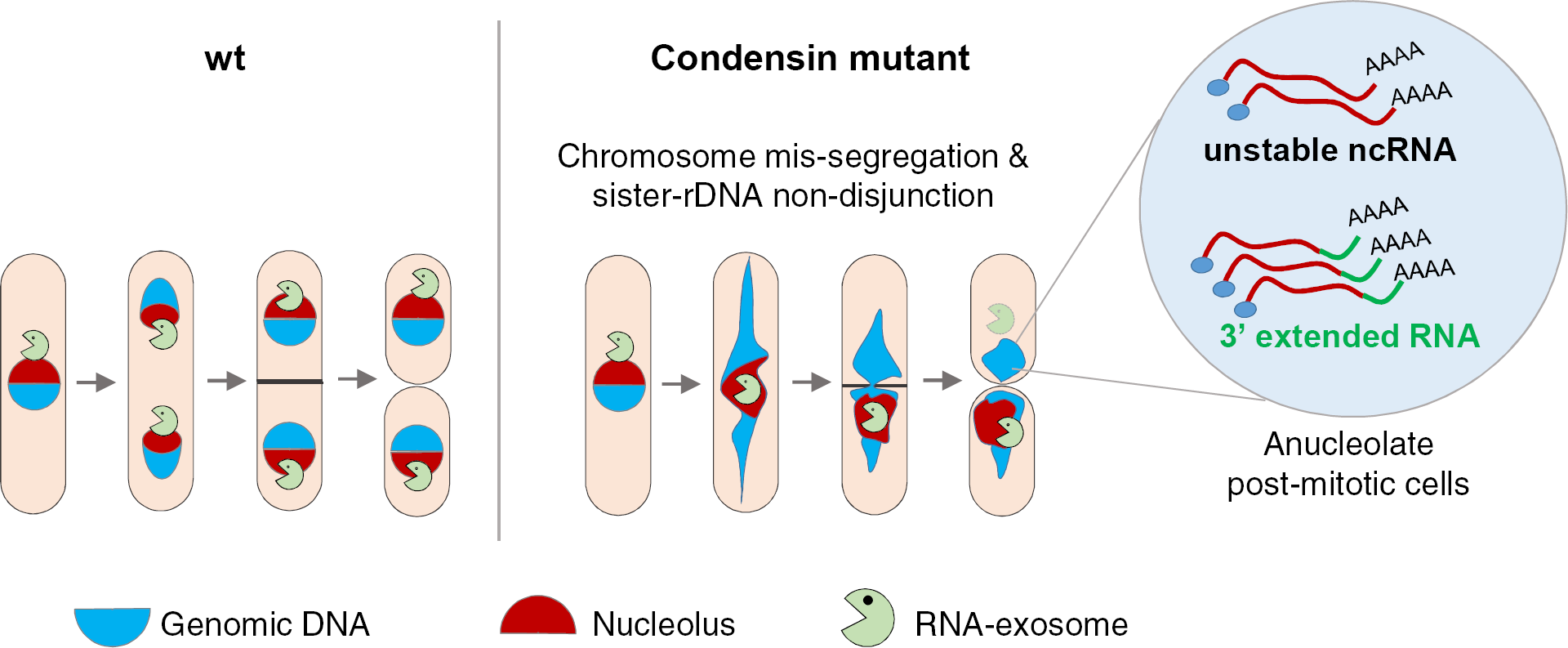
Condensin deficiency impinges upon gene expression by promoting accurate chromosome segregation throughout mitosis. In wild-type fission yeast cells, Rrp6 and Dis3, the catalytic subunits of the RNA-exosome, are enriched in the nucleolus and most likely co-segregate with the bulk of the rDNA during anaphase. In condensin mutant cells, chromosomes fail to properly segregate during anaphase. Chromatin bridges are formed and entangled sister-rDNA copies fail to separate. The non-disjunction of the rDNA upon mitotic exit leads to the production of anucleolate daughter cells deprived of Rrp6 and Dis3, which start to accumulate RNA molecules that are normally actively degraded by the RNA-exosome, such as unstable ncRNA and 3’-extended RNA.

That the bulk of the nucleolar RNA-exosome might co-segregate with the rDNA during mitosis has some consequences in the fields of gene silencing and RNA export. Condensin has been implicated in gene silencing at pericentric heterochromatin in fission yeast (He et al., 2016). However, since Rrp6 is known to take part in the turnover of heterochromatic transcripts (Buhler et al., 2007), it will be important to assess to which extent chromosome missegregation during anaphase might account for the alleviation of silencing observed in condensin-mutant cells. Likewise, genetic screens for factors involved in the nuclear export of polyadenylated mRNAs have led to the isolation of various unrelated mutations that all generate anucleolate cells, including mutations in condensin in budding yeast (Ideue et al., 2004; Paul and Montpetit, 2016). This has raised the idea that the nucleolus takes part in the nuclear export of mRNAs. However, since we observed no nuclear retention of *cct2* mRNA in anucleolate condensin-mutant cells, and since non-coding RNA are heavily polyadenylated in fission yeast (Zhou et al., 2015), we propose that the accumulation of polyadenylated transcripts in anucleolate cells might reflect an impaired processing of non-coding RNAs by the RNA-exosome rather than a defective export. The finding that Dis3 or Rrp6 loss-of-function increases the concentration of polyadenylated transcripts in the nucleolus of budding yeast cells is in good agreement with this conclusion (Paul and Montpetit, 2016).

Additional mechanisms linked to chromosome instability and the genesis of aberrant karyotypes in post-mitotic cells most likely contribute to modify the transcriptome when condensin I or II is defective. The slight up-regulation of histone genes that we observed in *cut14-208* mutant cells has previously been attributed to an indirect stabilization of the transcription factor Ams2 during defective anaphase (Kim et al., 2016). More broadly, aneuploidy itself is known to change the transcriptome, with a recurrent increased expression of genes involved in the response to stress and a down-regulation of cell-cycle and proliferation genes (Sheltzer et al., 2012). Premalignant murine T-cells defective for condensin II exhibit a transcriptome evocative of a stress response to aneuploidy (Woodward et al., 2016). Similarly, the depletion of Smc2, and hence the reduction of both condensin I and II, in a human neuroblastoma cell line modifies the expression of a large number of genes implicated mainly in cell cycle progression or DNA damage response (Murakami-Tonami et al., 2014). However, we did not observed such a clear transcriptomic signature in *cut14-208* mutant cells, suggesting either that aneuploidy per se, i.e. the gain or loss of entire chromosomes, is not a prevalent condition when condensin is impaired in fission yeast, or that condensin-mediated aneuploidy is strongly counter-selected in asynchronous cultures, perhaps because most aneuploid cells are unviable or extremely unstable in fission yeast (Niwa and Yanagida, 1985).

Our finding that sister-rDNA almost systematically non-disjoin during anaphase in fission yeast condensin mutant cells is reminiscent of the severe missegregation of the rDNA observed in budding yeast and human cells deprived of functional condensin complexes (D’Ambrosio et al., 2008; Freeman et al., 2000; Samoshkin et al., 2012), and is consistent with the idea that decatenation of sister-rDNA during late anaphase relies upon condensin (D’Ambrosio et al., 2008). What remains puzzling, however, is the fact that chromatin bridges are almost systematically severed by the cytokinetic ring in *cut14-208* mutant cells, while the severing of the nucleolus is extremely infrequent (<10% of the cases). The persistence of the nucleolus in the axis of cleavage might mechanically hinder cytokinesis, or trigger a wait signal. Alternatively, condensin deficiency might cause DNA double-strand breaks at fragile sites located on the centromere-proximal border of the cluster of rDNA repeats, allowing the displacement of untangled sister-rDNA towards one pole of the mitotic spindle through a spring-relaxation effect. A focused role for condensin in organising a region proximal to the rDNA, and located on its centromeric side, has been reported in budding yeast (Schalbetter et al., 2017). In HeLa cells, the depletion of SMC2 induces DNA breaks predominantly in repetitive DNA, including the rDNA/Nucleolar Organising Regions (Samoshkin et al., 2012). Thus missegregation of the rDNA in fission yeast cells when condensin is impaired might reveal an evolutionarily-conserved acute dependency of repeated DNA elements upon condensin for their segregation and integrity. Given the prevalence of tandem repetitive DNA arrays in mammalian genomes (Warburton et al., 2008), it is tempting to speculate that condensin loss of function might have similar confounding impacts on the preservation of their structural integrity and proper expression.

Studies performed over the last 20 years have shown that the tri-dimensional organization of the genome influences gene expression, raising the key question of the role played by SMC complexes in the control of gene expression. A large number of studies have concluded in favour of a role for cohesin and condensin I and II in the control of gene expression (Dowen and Young, 2014; Merkenschlager and Nora, 2016), raising the idea that cohesin and condensins might collectively link gene expression to genome architecture. Although there are robust examples of cohesin-mediated regulation of cell-type-specific gene expression (Merkenschlager and Nora, 2016), for instance through enhancer-to-promoter interactions (Ing-Simmons et al., 2015; Kagey et al., 2010), a recent study has raised the idea that the involvement of cohesin in the maintenance of an established gene expression program might be less prominent than initially thought (Rao et al., 2017). Similarly, despite ample reports of cells defective for condensin I or II exhibiting changes in gene expression (Bhalla et al., 2002; Dowen et al., 2013; He et al., 2016; Iwasaki et al., 2010; Kranz et al., 2013; Li et al., 2015; Longworth et al., 2012; Lupo et al., 2001; Murakami-Tonami et al., 2014; Rawlings et al., 2011; Wang et al., 2016; Yuen et al., 2017), to the best of our knowledge, there has been thus far no clear case where the influence of condensin I or II on gene expression has been dissociated from a possible confounding effect of chromosome missegregation. By providing evidence that condensin plays no major direct role in the control of gene expression in fission and budding yeasts, and showing that condensin impinges on gene expression by preserving the stability of the genome during mitosis, our work challenges the concept of gene regulation as a collective property of SMC complexes, and should help better define future studies on the role of canonical condensins in gene expression in other organisms.

## Materials and Methods

### Media, molecular genetics and strains

Media and molecular genetics methods were as previously described (Moreno et al., 1991). Fission yeast cells were grown at 28°C in complete YES+A medium or in synthetic PMG medium. The *nmt1-dis3* chimerical gene was repressed by the addition of thiamine 60 μM final to the growth medium, as described (Lemay et al., 2014), followed by further cell culture for 12–15h. Strains used in this study are listed in Table S2.

### Cell cycle synchronization

Fission yeast cells were synchronized in early S phase at 28°C by the adjunction of hydroxyurea (HU) 15 mM final. G2/M arrest was achieved using the thermo-sensitive *cdc25-22* mutation or the analogue-sensitive *cdc2asM17* allele (Aoi et al., 2014), and G1 arrest using the thermo-sensitive *cdc10-129* mutation. Reversible prometaphase arrest was performed at 19°C using the cold-sensitive *nda3-KM311* mutation (Hiraoka et al., 1984). Metaphase arrest was achieved in PMG medium supplemented with thiamine 20 µM using the thiamine repressible *nmt41-slp1* gene (Petrova et al., 2013). Mitotic indexes were measured as the percentages of mitotic cells accumulating Cnd2-GFP in the nucleus (Sutani et al., 1999). Budding yeast cells were grown at 30°C in YEPD to mid-log phase, collected by filtration, washed with dH_2_O, and re-suspended at an OD_600_ of 0.30.4 in YEPD containing 3 µg/ml α-factor. After one hour, additional α-factor was added to 3 µg/ml. After another hour, an aliquot of cells was used for ChIP (G1 sample) and the remaining cells were collected by filtration, washed with dH_2_O and re-suspended in YEPD to release the cells from the G1 arrest. 30 min after and 60 min after the release, aliquots were collected for ChIP (S phase sample and G2 sample respectively). Budding yeast strains expressing TEV protease under the *GAL1* promoter were grown at 30°C in YEP medium containing 2% raffinose (YEP-R) to mid-log phase, collected by filtration, washed with dH_2_O, and re-suspended at an OD600 of 0.15 in YEP-R containing 3 µg/ml α-factor. After one hour, additional α-factor was added to 3 µg/ml. After 30 min, the cultures were split and TEV protease expression was induced in one half by addition of 2% galactose. After another 30 min, cells were collected by filtration, washed with dH_2_O and re-suspended in YEP-R (uninduced) or YEP-R with 2% galactose (YEP-RG; induced) with 3 µg/ml α-factor. Fresh α-factor was added to 3 µg/ml after another hour to all cultures and ChIP samples were collected one hour later.

### Microscopy

To quantify anucleolate cells, cells expressing Gar2-mCherry and Psy1-GFP fusion proteins were fixed with cold methanol and DNA was stained with 4′, 6-diamidino-2-phenylindole (DAPI) at 0,5 μg/ml in PEM buffer (100 mM PIPES, 1 mM EGTA, 1mM MgSO4, pH 6,9). Gar2-mCherry and Psy1-GFP were directly observed under the fluorescent microscope. Rad22-GFP foci were analysed on cells fixed with iced-cold methanol and stained with DAPI. Cytological analysis of the CUT phenotype was performed as described (Hagan, 2016) except that cells were fixed with cold methanol and stained with Hoechst 33342 (20 μg/ml). Immunofluorescence was performed as described (Robellet et al., 2014), with the following modifications. Cells shifted at 36°C for 2h30 min were fixed with ice-cold ethanol and stored at 4°C. 2×10^7^ cells were washed in PEMS (PEM +1.2 M Sorbitol) and digested with Zymolyase 100T (0.4 mg/ml in PEMS) for 30 min at 37°C. Images were processed and quantified using ImageJ with automated background subtraction.

### FACScan

2×10^6^ fission yeast cells were fixed with ethanol 70% (v/v), washed in sodium citrate (50 mM pH 7) and digested with RNase A (100 µg/ml) (Merck). Cells were briefly sonicated and stained with 1 μM Sytox Green (ThermoFischer Scientific). DNA content was quantified on a FACSCALIBUR cytometer using CellQuest Pro software (BD Biosciences). Raw data were analyzed with FlowJo software (BD biosciences). FACScan analysis of budding yeast cells was performed as previously described after staining DNA with either propidium iodide (Fig. 4; Cuylen et al., 2011) or SYBR green I (Fig. S3; Cuylen et al., 2013).

### Chromatin immunoprecipitation and quantitative PCR

ChIPs for fission yeast cells were performed as described (Vanoosthuyse et al., 2014). 2×10^8^ cells were fixed with 1% formaldehyde at 36°C for 5 min and then 19°C for 25 min, washed with PBS and lysed using acid-washed glass beads in a Precellys homogenizer. Chromatin was sheared into 300-to 900-bp fragments by sonication using a Diagenode bioruptor. Sheared chromatin was split in two equivalent fractions subjected to parallel immunoprecipitations using magnetic Dynabeads coated with the appropriate antibody. Total and immunoprecipitated DNA was purified using the NucleoSpin PCR clean-up kit (Macherey-Nagel). DNA was analysed on a Rotor-Gene PCR cycler using QuantiFast SYBR Green mix. ChIP-qPCR experiments for budding yeast cells were performed as described previously (Cuylen et al., 2011). In brief, aliquots of 42 ml culture with an OD_600_ of 0.6 were fixed in 3% formaldehyde at 16°C. Chromatin was sonicated to an average length of 500 bp using a Diagenode bioruptor. For anti-HA immunoprecipitation, 50 µl protein G dynabeads and 1.5 µl 16B12 antibody (anti-HA.11, Covance) were used. For anti-PK immunoprecipitation, 50 µl protein A dynabeads and 2 µl anti-PK (V5) tag antibody (Abd Serotec MCA1360) were used. Purified DNA was analysed with an ABI 7500 real-time PCR system (Applied Biosystems) using rDNA-specific primers. Primers are listed in Table S3.

### Chromosome spreads

Budding yeast cells were grown at 30°C in YEPD to mid-log phase, collected by filtration, washed with dH_2_O, and re-suspended at an OD_600_ of 0.2 in YEPD containing 3 µg/ml α-factor. After one hour, additional α-factor was added to 3 µg/ml. After another hour, an aliquot of cells was used for chromosome spreading (G1 sample) and the remaining cells were collected by filtration, washed with dH_2_O and re-suspended into YEPD to release the cells from the G1 arrest. Aliquots were collected for chromosome spreading 30 min and 60 min after the release (S and G2 phase samples, respectively). Chromosome spreads were prepared as described previously (Cuylen et al., 2011) and stained for Brn1-HA_6_ with 16B12 (anti-HA.11, Covance, 1:500) and Alexa Fluor 594–labelled anti-mouse IgG (Invitrogen, 1:600) antibodies and for DNA with DAPI. Images were recorded on a DeltaVision Spectris Restoration microscope (Applied Precision) with a 100×, NA 1.35 oil immersion objective.

### Total RNA extraction and RT-qPCR

Total RNA was extracted from 2×10^8^ fission yeast cells by standard hot-phenol method. 1 µg of total RNA was reverse-transcribed using Superscript III (Life Technologies) following the manufacturer’s instructions, using random hexamers in the presence or absence of Reverse Transcriptase (RT). cDNAs were quantified by real time qPCR on a Rotor-Gene PCR cycler using QuantiFast SYBR Green mix. The absence of PCR product in minus RT samples has been verified for all RT-qPCR shown in this publication. Primers are listed in Table S3.

### RNase digestion and Northern blot

For RNase H digestion, total RNA was hybridized with a DNA oligonucleotide complementary to a sequence located at the 5’ end of the *hsp9* mRNA and digested with RNase H (Roche) following the manufacturer’s instructions. For Northern blotting, total or RNase H-digested RNA was resolved on 1% agarose gel supplemented with formaldehyde 0.7% (v/v), transferred onto Hybond-XL nylon membranes (Amersham) and cross-linked. Pre-hybridization and overnight hybridization were carried out in ULTRAhyb buffer (Ambion) at 68°C. Strand-specific RNA probes were generated by in vitro transcription using the T7 riboprobe system (Ambion) and internally labelled with [α-32P]-UTP. Membranes were quickly washed with 2X SSC and 0,1% SDS, 10 min in 2X SSC and 0,1% SDS, and 3 times in 0,2X SSC and 0,1% SDS. Blots were imaged with Typhoon 8600 instrument (Molecular Dynamics) and quantified with ImageQuant TL (GE Healthcare).

### Single molecule RNA Fluorescence In Situ Hybridization (smFISH), imaging and quantification

smFISH was performed on formaldehyde-fixed cells, as described (Keifenheim et al., 2017). Probes were designed and synthetized by Biosearch Technologies (Petaluma, CA). The *mug93as* and *cct2* probes were labelled with Quasar 670. Probe sequences are listed in Table S3. Cells were imaged on a Leica TCS Sp8, using a 63x/1.40 oil objective, Optical z sections were acquired (z-step size 0.3 microns) for each scan to cover the depth of the cells. For image analysis and quantification, cell boundaries were outlined manually and RNA spots quantified using FISH-quant package implemented in MATLAB, as described (Mueller et al., 2013). The FISH-quant detection technical error was estimated at 6-7% by quantifying mRNA numbers using two sets of probes covering the 5’ half or the 3’ half of the *rpb1* mRNA and labelled with different dyes.

### RNA-seq and analysis

RNA-seq was performed on biological triplicates. Total RNA was extracted from 2×10^8^ yeast cells by standard hot-phenol method. RNA quality was determined using TapeStation (Agilent) with RINe score >9. Ribosomal RNA was removed by treating 2 µg of total RNA with the Ribo-Zero Gold rRNA Removal Kit (Yeast) (MRZY1324, Illumina, Paris, France). RNA-seq libraries were prepared using TruSeq Stranded kit. Sequencing was performed on Illumina Hiseq 4000, with single-end reads of 50 nt in length. Total number of reads per sample ranged from 59 million to 93 million. For in-silico analyses, we used the version 2.30 of the *S. pombe* genome in fasta format and the corresponding gff3 annotations downloaded from the ebi website (2017/03/22). Scripts are available in the following git repository: https://github.com/LBMC/readthroughpombe. The RNA-seq reads were processed with cutadapt to remove adaptors and further trimmed with UrQt (--t 20) based on their quality. After quality control with fastqc, we built the reverse complement of the reads in the fastq files using seqkit, indexed the genome (-build) and mapped the trimmed fastq files (--very-sensitive) using Bowtie2. Mapping output files were saved in bam format with samtools (view-Sb).

To detect read-through events, we searched for sections of reads located immediately downstream of the 3’ ends of annotated transcription units, on the same DNA strand, and within gene-free intergenic regions. First, broad peaks of RNA-seq reads were identified using the peak caller Music (v6613c53). Bam files were sorted and filtered as forward or reverse with samtools (view -hb -F 0×10 or view -hb -f 0×10). The gff annotation file was converted to bed with convert2bed from bedtools, the genome indexed with samtools (faidx) and the annotation file split into forward and reverse with bedtools (complement). Next, reads corresponding to annotated transcripts were removed using samtools view (-hb bams -L annotation). Subtracted bam files were sorted with samtools sort, and RNA-seq peaks located outside annotated transcripts were detected with Music (-get_per_win_p_vals_vs_FC -begin_l 50 -end_l 500 -step 1.1 -l_mapp 50 -l_frag 363 -q_val 1 -l_p 0). The resulting annotation was further filtered by removing peaks whose starting positions were located more than 2 read-length away from the nearest 3’ends. Only peaks detected in at least 2 out of 3 biological replicates were considered. Read-through detection was performed independently for the *rrp6Δ* and *cut14-208* mutants. To achieve comparable examination between *rrp6Δ* and *cut14-208* for reads quantification, we merged their respective read-through annotations and sorted them with bedtools (sort) and extracted the forward and reverse reads from the bam files with bedtools (bamtofastq). We generated the list of transcript sequences from the genome and the annotation with bedtools (getfasta -s). The transcript sequences were then indexed with kallisto (index-k 31 --make-unique) and the quantification achieved with kallisto (quant –single -l 363.4 -s 85.53354). Quantifications were performed separately on the transcript alone and on the transcript plus read-through annotation.

Quantifications in mutant strains compared to wt were performed using the package DESeq2 with R (v3.4.4). For differential expression analyses, we tested for log2 fold change superior to 0.5 or inferior to −0.5. For read-through events, we tested for a log2 fold change superior to 0 compared to the wild type to declare the read-through present in the mutant background. For all analyses, we selected p-values with an FDR <= 0.05. The package ggplot2 (v2.2.1) was used for graphics.

### High-resolution tiling arrays

For transcriptome analysis in G1 phase, *Δbar1* cells were grown at 30°C in YEP-R to mid-log phase, collected by filtration, washed with dH_2_O, and re-suspended at an OD_600_ of 0.3–0.4 in YEP-R containing 3 µg/ml α-factor. After 1.5 h, galactose was added to 2% and, after another 2.5 h, 100 ml cells of OD_600_= 0.6–0.8 were harvested by centrifugation at room temperature for RNA isolation. For transcriptome analysis during G2 phase, cells were grown at 30°C in YEP-R to mid-log phase, collected by filtration, washed with dH_2_O, and re-suspended at an OD_600_ of 0.3–0.4 in YEP-R containing 3 µg/ml α-factor. After one hour, additional α-factor was added to 3 µg/ml and 30 min later galactose was added to 2%. After another 30 min, cells were collected by filtration, washed with dH_2_O and re-suspended in YEP-RG with 3 µg/ml α-factor. Fresh α-factor was added to 3 µg/ml after one hour. After another hour, cells were collected by filtration, washed with dH_2_O and re-suspended in YEP-RG with 10 µg/ml nocodazole. 1.5 h after the release from G1 phase, 100 ml culture with OD_600_ of 1.0 were harvested by centrifugation at room temperature for RNA isolation. Samples were collected for FACScan analysis at the indicated time points.

High-resolution tiling arrays were performed and analysed as described (Xu et al., 2009). In brief, RNA isolated from yeast cells was reverse transcribed to cDNA with a mixture of random hexamers and oligo-dT primers, labelled and hybridized to tiled Affymetrix arrays of the budding yeast genome (S288c Genome Chip, http://www-sequence.stanford.edu/s288c/1lq.html). Transcripts that were two-fold or more up-or downregulated are listed in Table S5.

### Accession number

RNA-seq data are accessible from the Gene Expression Omnibus (GEO) database under the accession number GSE112281.

### Antibodies

Antibodies used in this study are listed in Table S4.

## Acknowledgements

We thank Vincent Vanoosthuyse and André Verdel for fruitful discussions, and V. Vanoosthuyse for critical reading of the manuscript. We are grateful to François Bachant, Kim Nasmyth, Chris Norbury, Jean-Paul Javerzat, Masayuki Yamamoto and the Yeast Genetic Resource Center of the National BioResource Project–Japan for strains, and to Keith Gull for the TAT1 antibody. We thank the Pôle Scientifique de Modélisation Numérique (PSMN) of ENS-Lyon and the Biocomputing Pole of the LBMC for in-silico analyses. We acknowledge the help of Hélène Polveche and Lorraine Soudade during the initial phase of bioinformatics analyses. Clémence Hocquet is supported by a PhD studentship from the Ministère de l’Enseignement Supérieur et de la Recherche and from the Fondation pour la Recherche Médicale (grant FDT20170437039). Xavier Robellet is supported by a post-doctoral fellowship from the ANR. Research in the Bernard lab is supported by the CNRS, the ANR (grant ANR-15-CE12-0002-01), the Fondation ARC pour la Recherche sur le Cancer (grant PJA 20151203343) and the Comité du Rhône de la Ligue Nationale contre le Cancer. This research in the Marguerat lab was supported by the UK Medical Research Council. Research in the Haering and Steimetz labs is supported by EMBL.

## Competing interests

The authors declare no financial and non-financial competing interest

---

**Supplemental Figure S1 - related to Figure 1.**
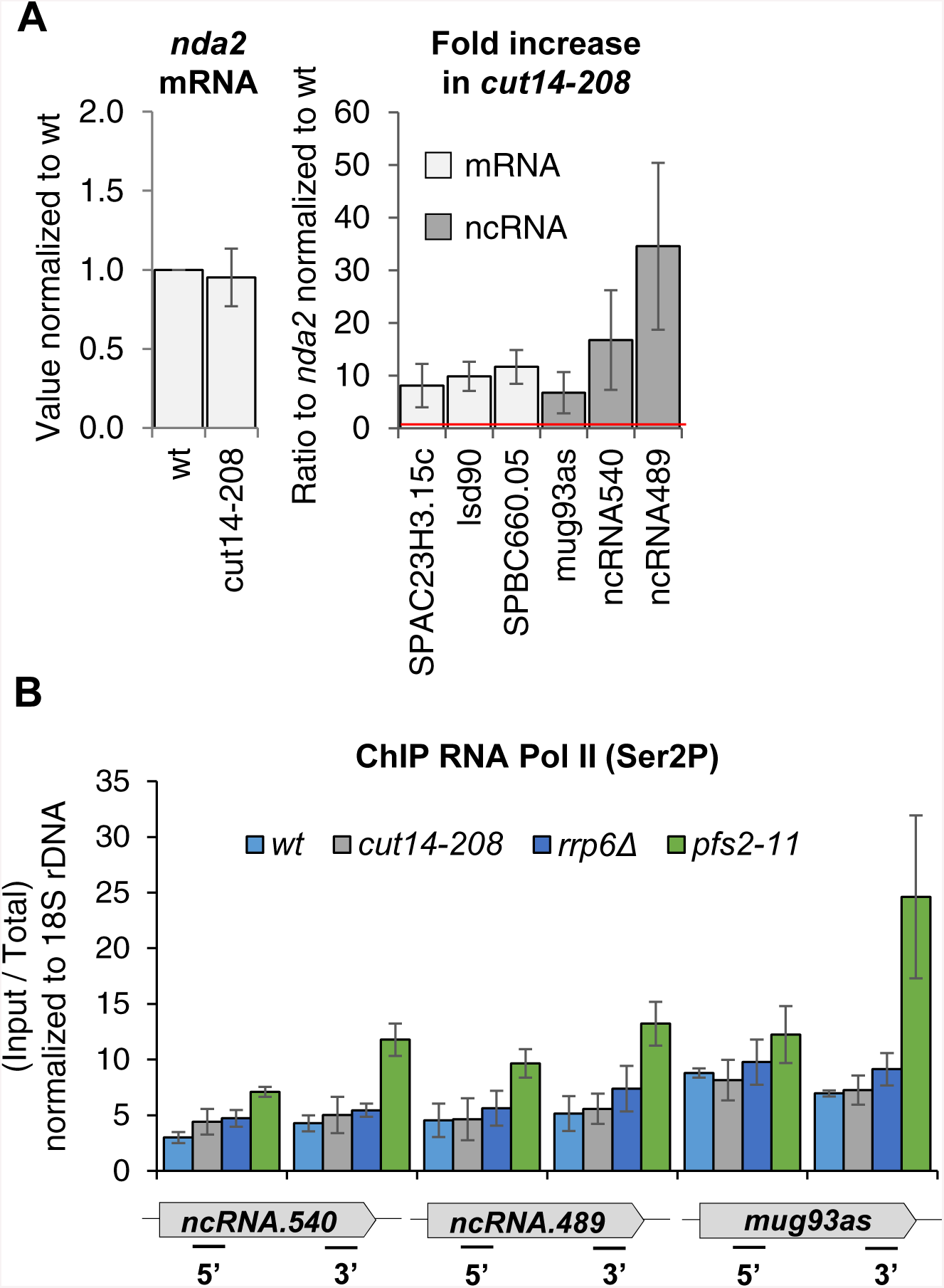
The condensin loss-of-function mutant *cut14-208* accumulates exosome-sensitive transcripts. **A.** RT-qPCR validation of up-regulated RNA in *cut14-208* mutant cells. Total RNA extracted from wild-type and *cut14-208* cycling cells grown for 2.5 h at the restrictive temperature of 36°C was reverse-transcribed in the presence or absence of RT and the cDNA quantified by qPCR. Shown are averages ± SDs measured from n = 3 biological replicates. **B.** Pol II occupancy measured by ChIP. Cells of indicated genotypes were grown at the restrictive temperature of 36°C for 2.5 hours and processed for ChIP against RNA Pol II phosphorylated on serine 2 of the CTD. The transcription termination mutant *pfs2-11* was used as a control for Pol II accumulation in the 3’ ends of genes (Wang et al., 2005). Show are averages ± SDs calculates from 6 ChIPs performed on biological triplicates.

**Supplemental Figure S2 - related to Figure 2.**
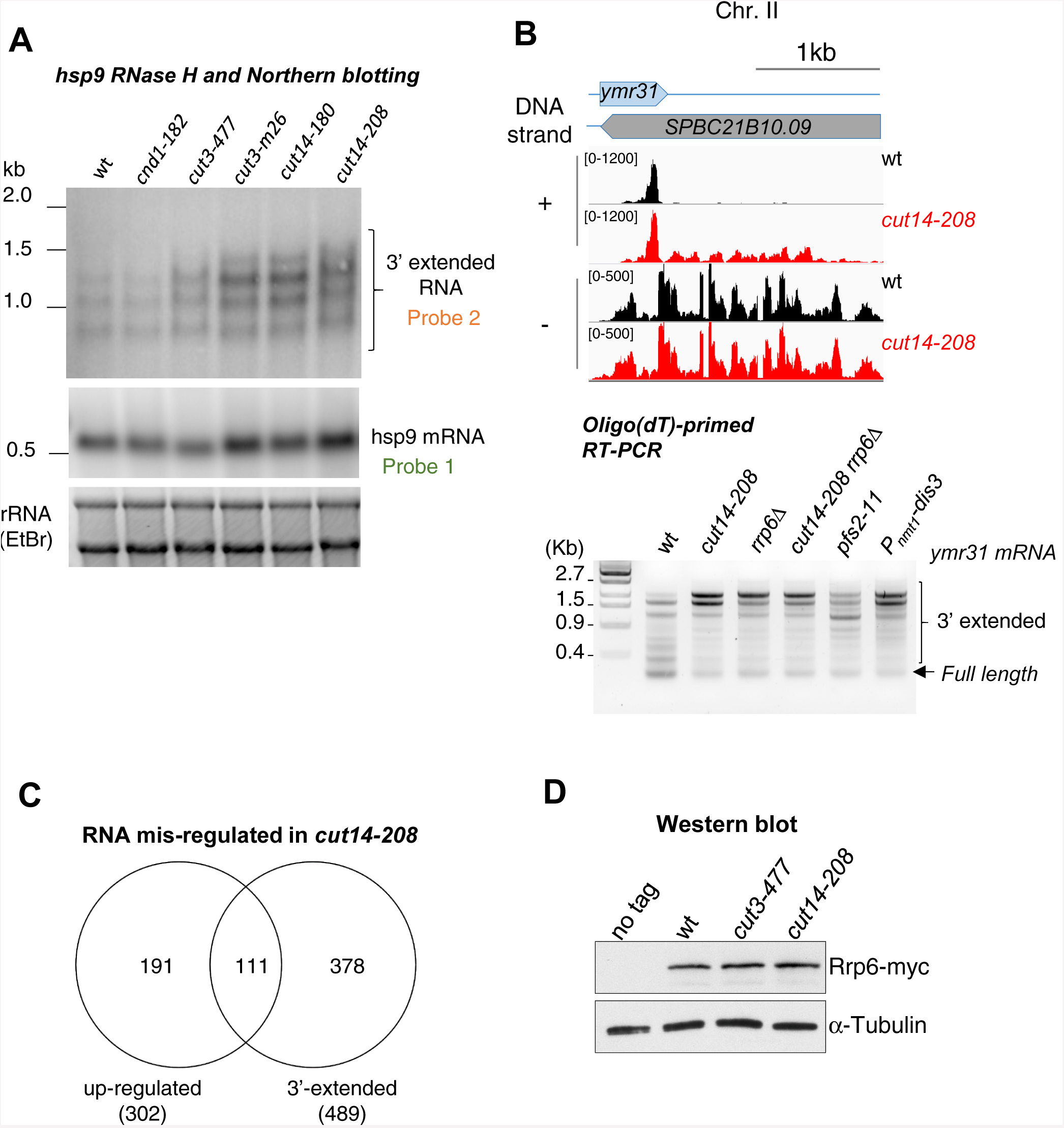
The condensin loss-of-function mutant *cut14-208* accumulates 3’-extended read-through transcripts. **A.** 3’-extended *hsp9* RNA detected by RNase H digestion and Northern blotting in condensin mutant cells. Indicated strains were grown at 36°C for 2.5 hours. Total RNA was digested by RNase H in the presence of a DNA oligonucleotide complementary to the 5’end of *hsp9* mRNA. Cleaved products were revealed by a probe hybridizing downstream of the transcription termination site of *hsp9* (probe 2 in Fig. 2A), or within the coding sequence (probe 1 in Fig. 2A). rDNA stained with EtBr served as loading control. **B.** Polyadenylated RNAs detected by oligo(dT)-primed RT-PCR. Cells were grown at 36°C for 2.5 hours in PMG supplemented with 20 µM thiamine to repress *nmt1-dis3*. Total RNA was reverse transcribed using oligo(dT) primers in the presence or absence of RT. cDNA were amplified by 25 cycles of PCR using oligo(dT) and gene specific primers. PCR products were separated on an agarose gel and stained with EtBr. Minus RT reactions produced no signal. **C.** Venn diagram showing the overlap between the two sets of increased RNAs and read-through RNAs in *cut14-208*. **D.** Steady state level of Rrp6. Indicated strains were grown at 36°C for 2.5 hours, total proteins were extracted and probed with an anti-myc antibody. Alpha-tubulin served as loading control.

**Figure S3 - related to Figure 4.**
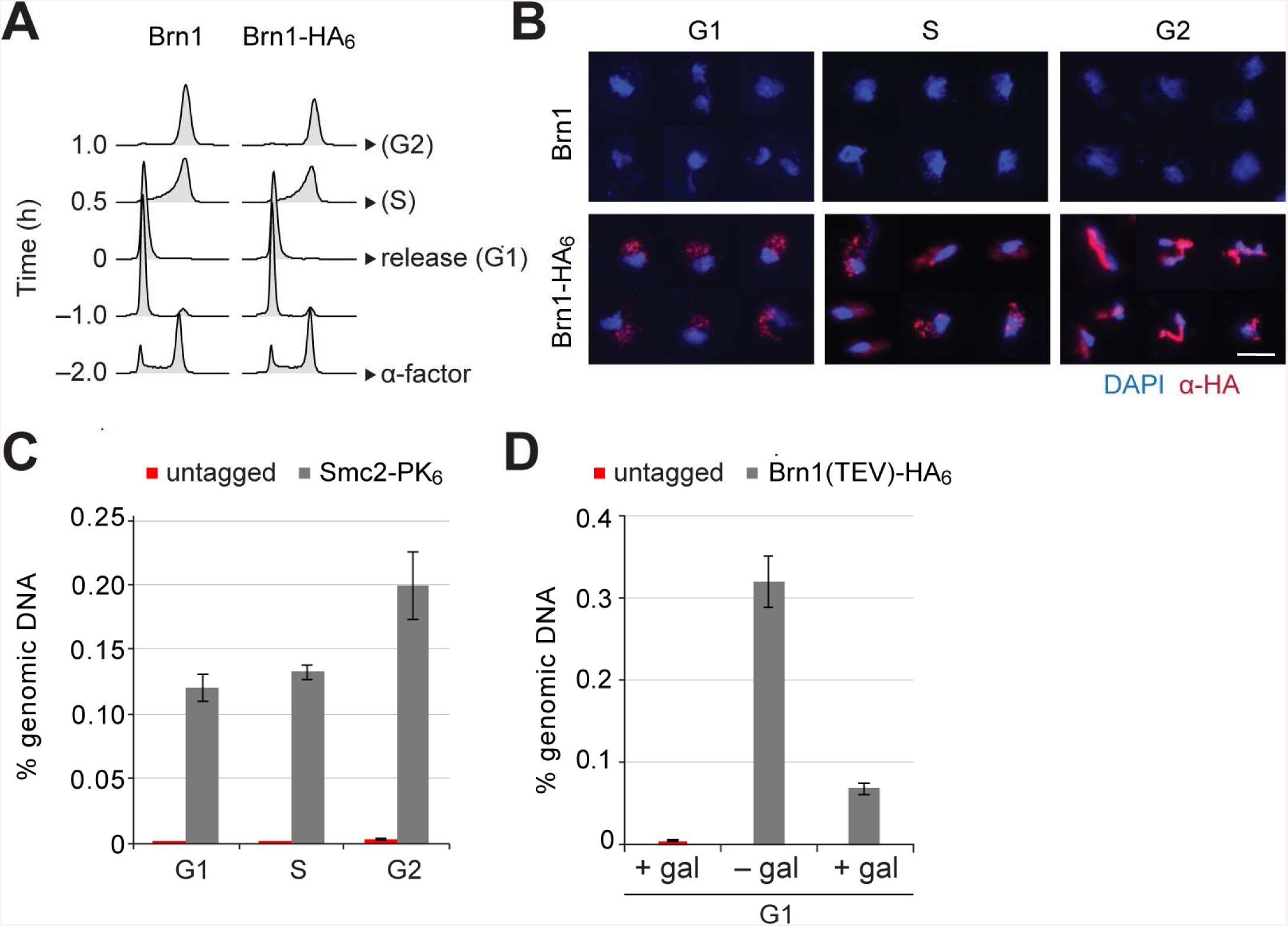
Condensin binds to budding yeast chromosomes throughout the cell cycle and is released by TEV cleavage of its kleisin subunit. **A**. Cells were synchronized to G1 phase by α-factor and released (strains C1 and C1584). Samples for chromosome spreads were taken prior to the release, 0.5 and 1 h after the release in order to yield G1, S and G2 phase samples, as indicated. Cell cycle synchronization was scored by FACScan analysis of cellular DNA content. **B.** Condensin levels on chromosomes of cells in **A** were measured by immunofluorescence of chromosome spreads (red, anti-HA; blue DAPI). Scale bar: 5 µm. **C.** Cells were synchronized to G1 phase by α-factor (strains C1 and C1597) and released as in **A**. Samples for ChIP were taken prior to the release, or 0.5 or 1 h after the release for G1, S and G2 phase samples, respectively. Condensin levels on chromosomes were measured by anti-PK ChIP at the rDNA locus followed by quantitative PCR. **D.** TEV protease expression was induced by addition of galactose (+ gal) in cells synchronized in G1 phase by α-factor (strains C1039 and C2455). A control sample was not induced (– gal). Condensin levels on chromosomes were measured by anti-HA ChIP followed by quantitative PCR at the rDNA locus.

**Supplemental Figure S4 - related to Figure 5.**
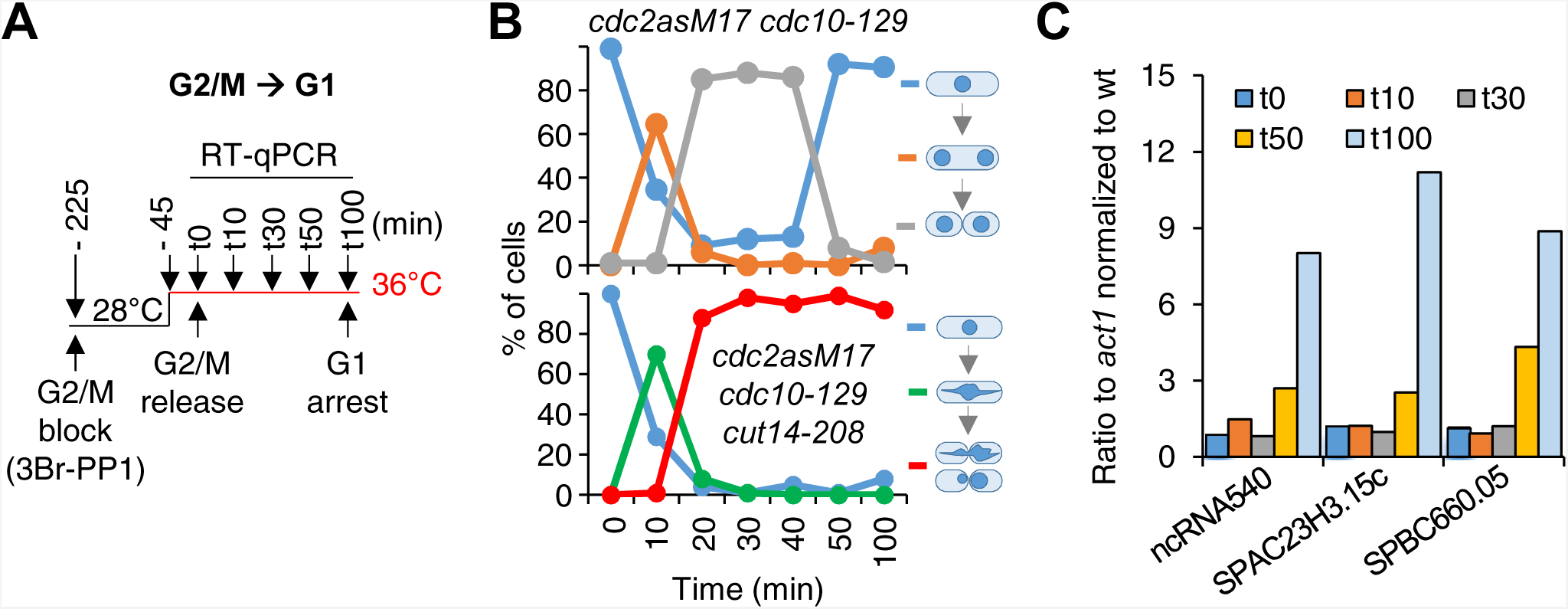
Defective mitosis underlies deregulated gene expression in the fission yeast *cut14-208* condensin mutant. **A-C.** Gene expression was assessed in synchronized *cut14-208* cells progressing through mitosis. **A.** Cells expressing the analogue-sensitive Cdc2asM17 kinase were arrested at the G2/M transition in the presence of 3-Br-PP1 (2 µM), shifted at 36°C, released into mitosis and re-arrested in late G1 phase by the *cdc10-129* mutation. **B.** DNA content was assessed by FACScan analysis and mitotic progression by cytological observation. DNA was stained with DAPI and the septum with calcofluor. The frequencies of chromatin bridges (red line) and chromosome cutting by the septum (green line) are shown. **C.** Total RNA extracted from wild-type and *cut14-208* cells shown in **B** was reverse-transcribed in the presence or absence of RT and cDNA quantified by qPCR.

**Supplemental Figure S5 - related to Figure 5.**
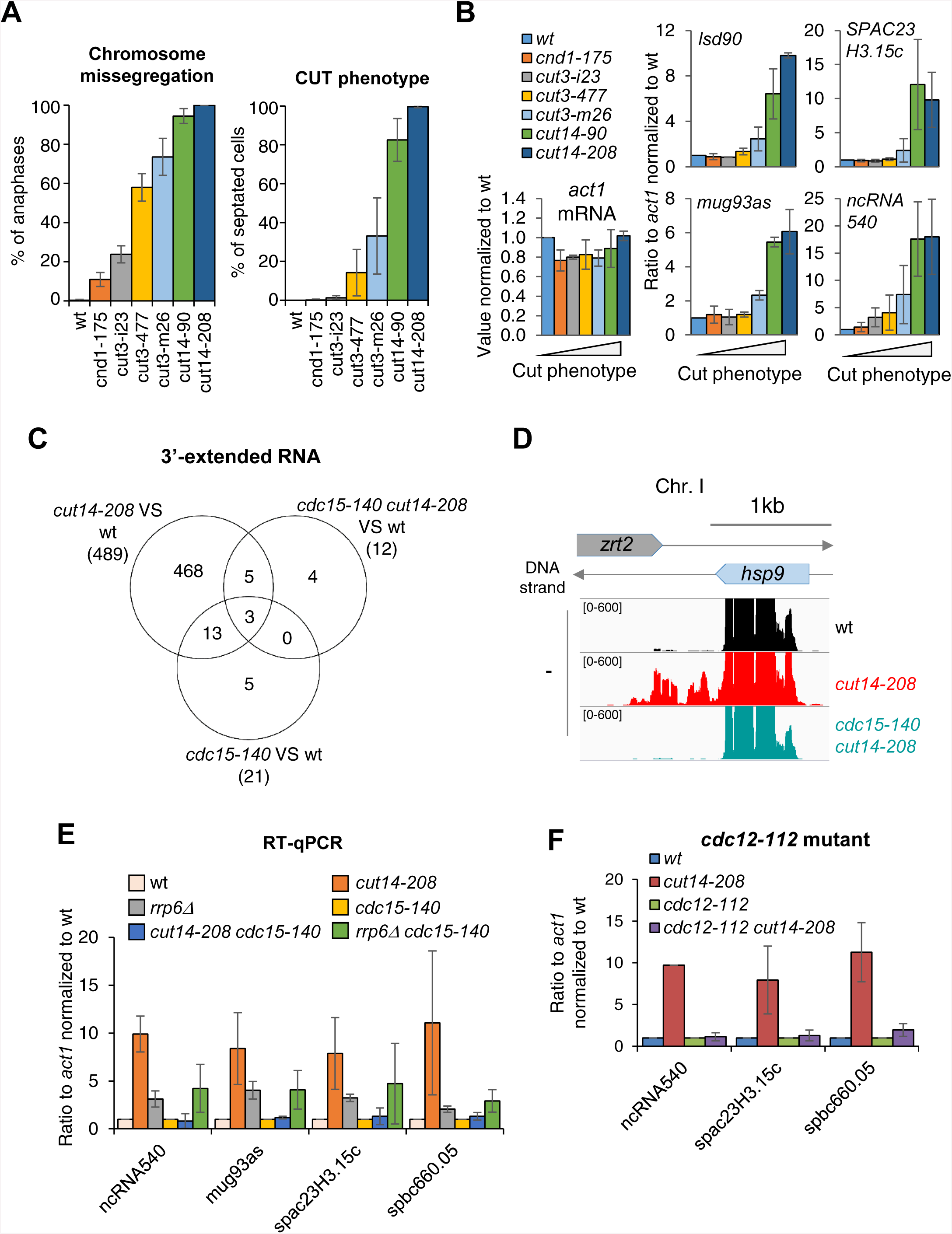
Defective mitosis underlies deregulated gene expression in the fission yeast *cut14-208* condensin mutant. **A-B.** The amplitude of increase in RNA levels correlates with the prevalence of chromosome severing. Indicated cells were grown at 36°C for 2.5 hours and processed for cytological analysis of chromosome segregation or RT-qPCR. **A.** Cells were fixed and processed for immunofluorescence against α-tubulin. DNA was stained with DAPI. Chromosome segregation was assessed in anaphase cells exhibiting a mitotic spindle ≥ 6 µm in length (n ≥ 100). CUT phenotype (chromosome severing by the septum) was assessed by staining DNA with Hoechst 3342 and the septum by calcofluor (n ≥ 100). Shown are averages ± SDs from biological triplicates. **B.** Total RNA extracted from cells grown at 36°C was reverse-transcribed in the presence or absence of RT and the cDNA quantified by qPCR. Shown are the averages ± SDs measured from biological triplicates. **C.** Venn diagram of 3-extended RNA detected by strand specific RNA-seq in indicated strains. **D.** RNAseq profiles of the *hsp9* gene showing the suppressive effect of *cdc15-140* on the accumulation of read-through transcripts caused by *cut14-208*. **E.** Increased RNA levels persist in the double mutant *cdc15-140 rrp6Δ.* Cells were grown at 36°C for 2.5 hours, total RNA was reverse-transcribed in the presence or absence of RT and cDNA quantified by qPCR. Shown are the averages ± SDs measured from biological triplicates. **F.** The mutations *cdc12-112* that prevents cytokinesis at the restrictive temperature restored normal RNA levels in a *cut14-208* genetic background. Cells were grown at 36°C for 2.5 hours, total RNA was reverse-transcribed in the presence or absence of RT and cDNA quantified by qPCR. Shown are the averages ± SDs measured from biological duplicates.

**Supplemental Figure S6 - related to Figure 6.**
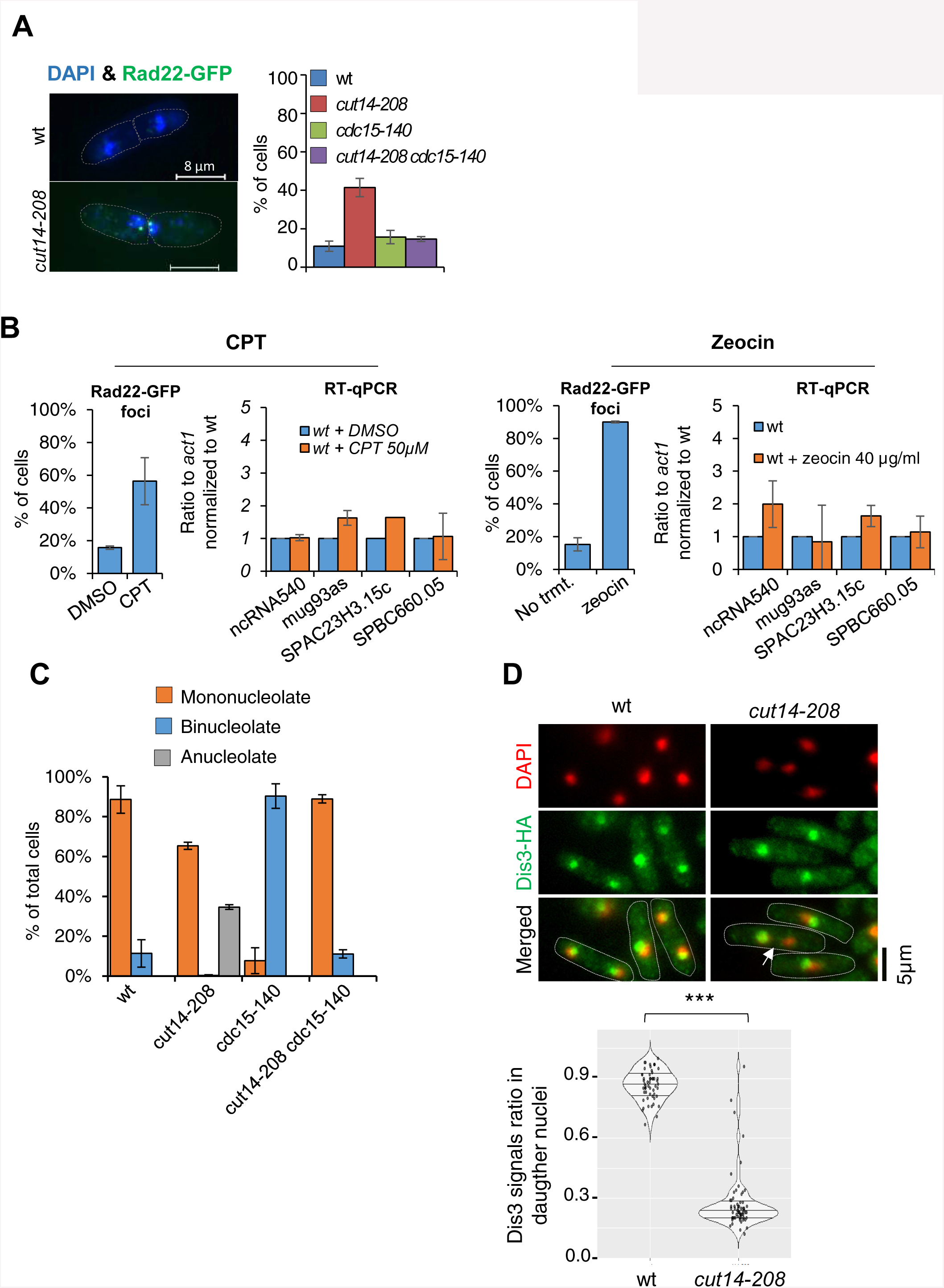
Condensin inactivation generates anucleolate daughter cells in fission yeast, which are depleted of the RNases Rrp6 and Dis3 and accumulate unstable RNA. **A.** *cut14-208* mutant cells accumulate DNA damage that can be suppressed by *cdc15-140*. Cells expressing Rad22-GFP were grown at 36°C for 2.5 hours, fixed and stained with DAPI. Cells exhibiting at least one Rad22-GFP focus were scored. Shown are the averages ± SDs calculated from biological triplicates with more than 100 cells per experiment. **B.** DNA damage does not increase RNA levels as *cut14-208*. Cells expressing Rad22-GFP were synchronised in prometaphase by the *nda3-KM311* mutation and released in mitosis in the presence of Camptothecin (CPT), or its vehicle DMSO, to induce DNA damage upon mitotic exit. Asynchronously growing cells expressing Rad22-GFP were treated with zeocin to induce DNA double-strand breaks. The prevalence of DNA damage was measured by scoring Rad22-GFP foci. RNA levels in cells experiencing DNA damage were assessed by RT-qPCR. Shown are averages ± SDs calculated from biological triplicates. **C.** The *cdc15-140* mutation prevents the production of anucleolate cells in a *cut14-208* genetic background. Cells expressing Gar2-GFP were grown at 36°C for 2.5 hours, fixed and stained with DAPI to score the percentage of anucleolate cells. Shown are the averages ± SDs calculated from biological triplicates with more than 100 cells per experiment. **D.** Dis3 is enriched in the nucleolus and segregates asymmetrically in *cut14-208* mutant cells (arrow). Indicated cells were grown at 36°C, fixed with ethanol and processed for immunofluorescence against Dis3-HA. DNA was stained with DAPI. Right panel shows the ratio of Dis3-HA signals measured within daughter nuclei in septated cells. *** indicates that the difference is statistically significant with p<0.001 by the Chi-square test.

**Table S1.** RNA levels of RNA-exosome and TRAMP components in *cut14-208* mutant cells

| Protein | Systematic gene name | Protein Complex | *cut14-208* vs wt |  |
| --- | --- | --- | --- | --- |
|  |  |  | P value adj | Log2FC |
| Rrp6 | SPAC1F3.01 | RNA-exosome | 1 | -0,24371 |
| Dis3 | SPBC26H8.10 | RNA-exosome | 1 | -0,13197 |
| Rrp41 | SPAC3G9.10c | RNA-exosome | 1 | -0,23751 |
| Rrp43 | SPBC17D1.03c | RNA-exosome | 1 | -0,10895 |
| Cti1/Rrp47 | SPCC1739.07 | RNA-exosome | 1 | 0,00185 |
| Csl4 | SPCC1840.11 | RNA-exosome | 1 | -0,13319 |
| Rrp4 | SPAC2F7.14c | RNA-exosome | 1 | -0,18844 |
| Rrp40 | SPAC22A12.12c | RNA-exosome | 1 | 0,02820 |
| Rrp46 | SPBC115.01c | RNA-exosome | 1 | -0,16979 |
| Mtr3 | SPBC211.08c | RNA-exosome | 1 | 0,03692 |
| Rrp42 | SPBC16G5.10 | RNA-exosome | 1 | -0,01230 |
| Rrp45 | SPCC757.08 | RNA-exosome | 1 | -0,09132 |
| cid14 | SPAC12G12.13c | TRAMP | 1 | -0,17385 |
| Air1 | SPBP35G2.08c | TRAMP | 1 | -0,32050 |
| Mtr4 | SPAC6F12.16c | TRAMP | 1 | -0,02359 |
| RNA helicase | SPAC17H9.02 | TRAMP | 1 | -0,22135 |

**Table S2.**
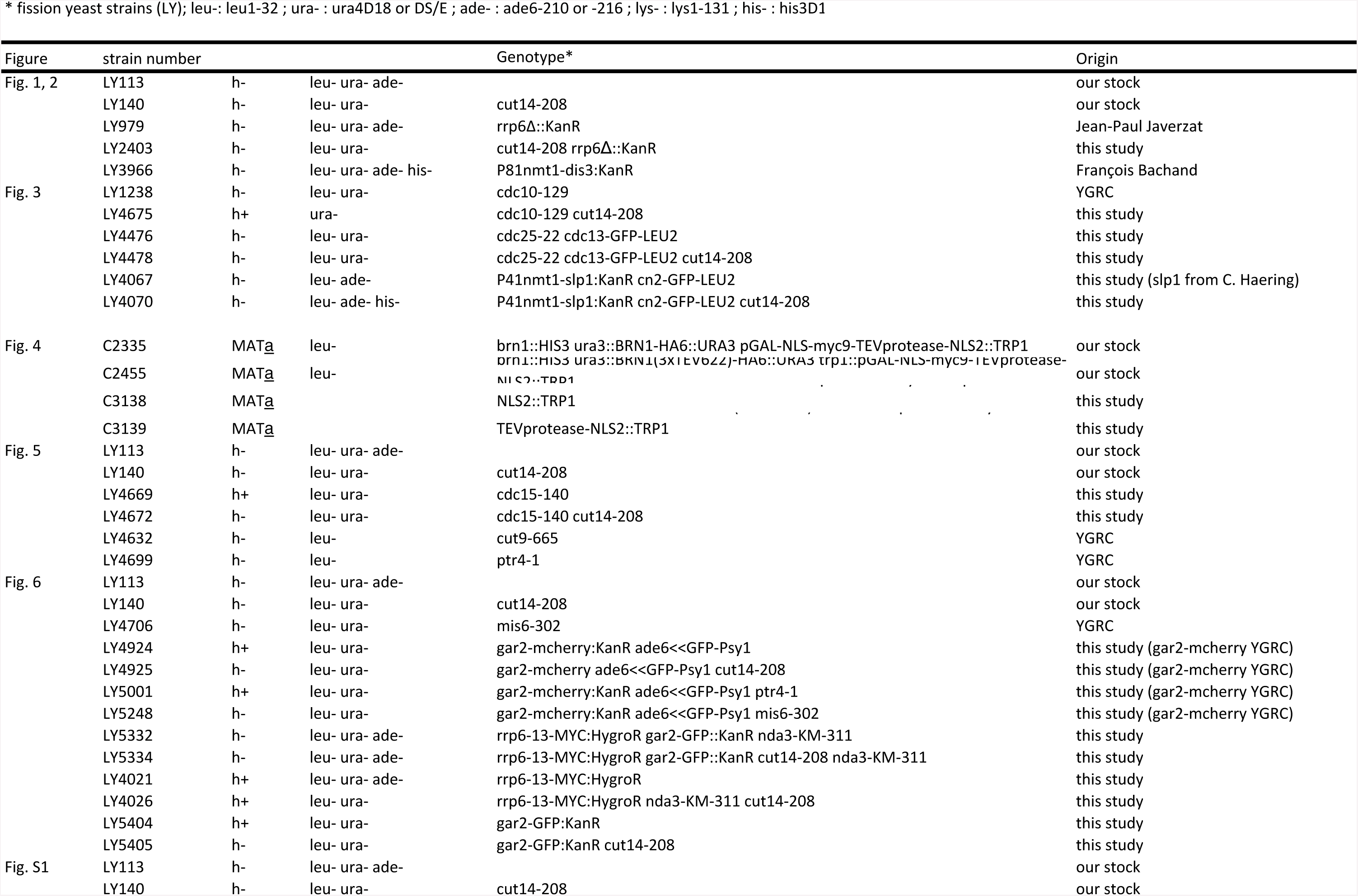
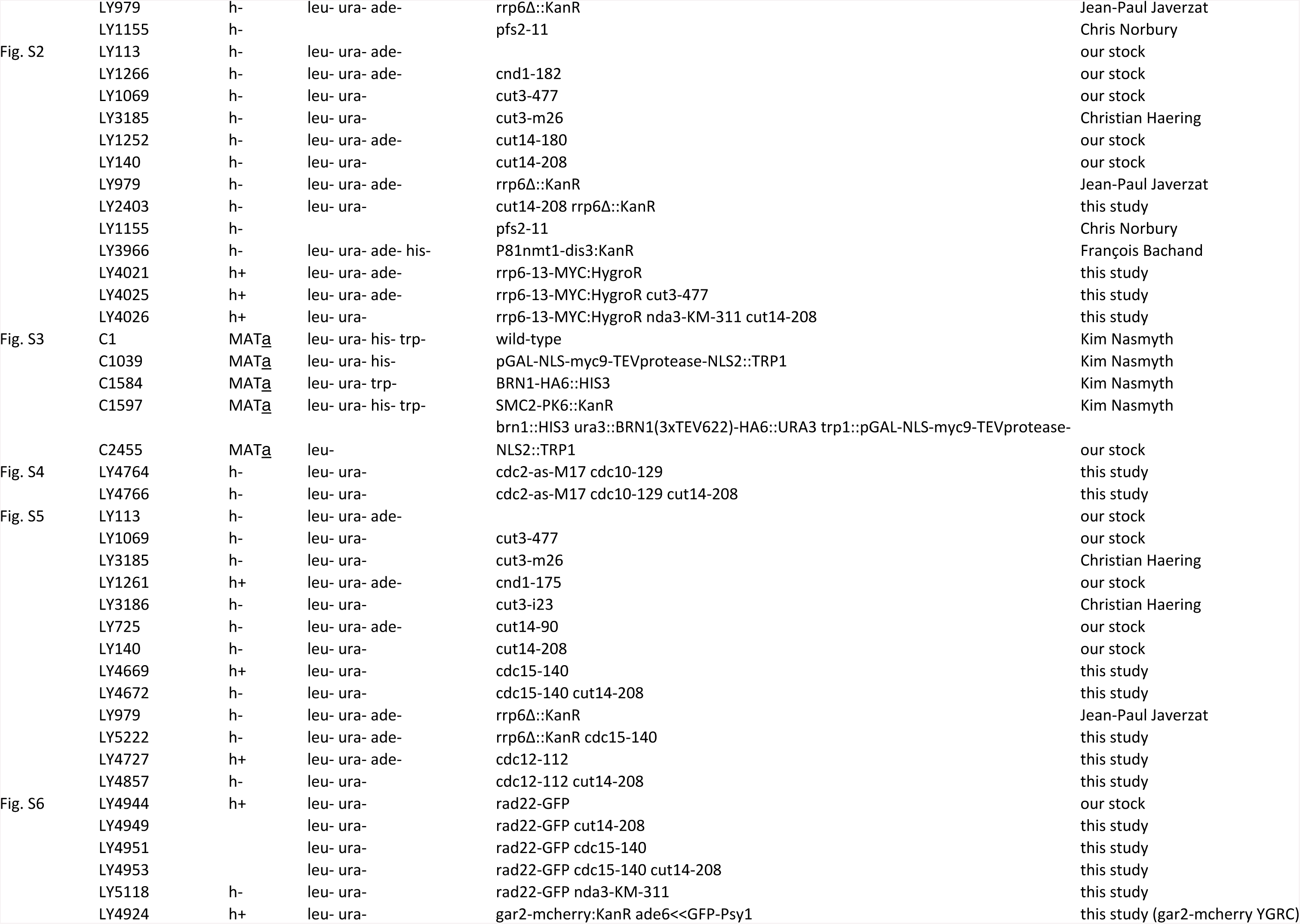
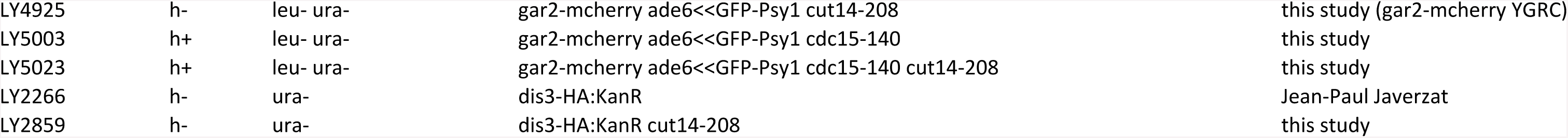
Yeast strains used in this study

**Appendix Tables S3:**
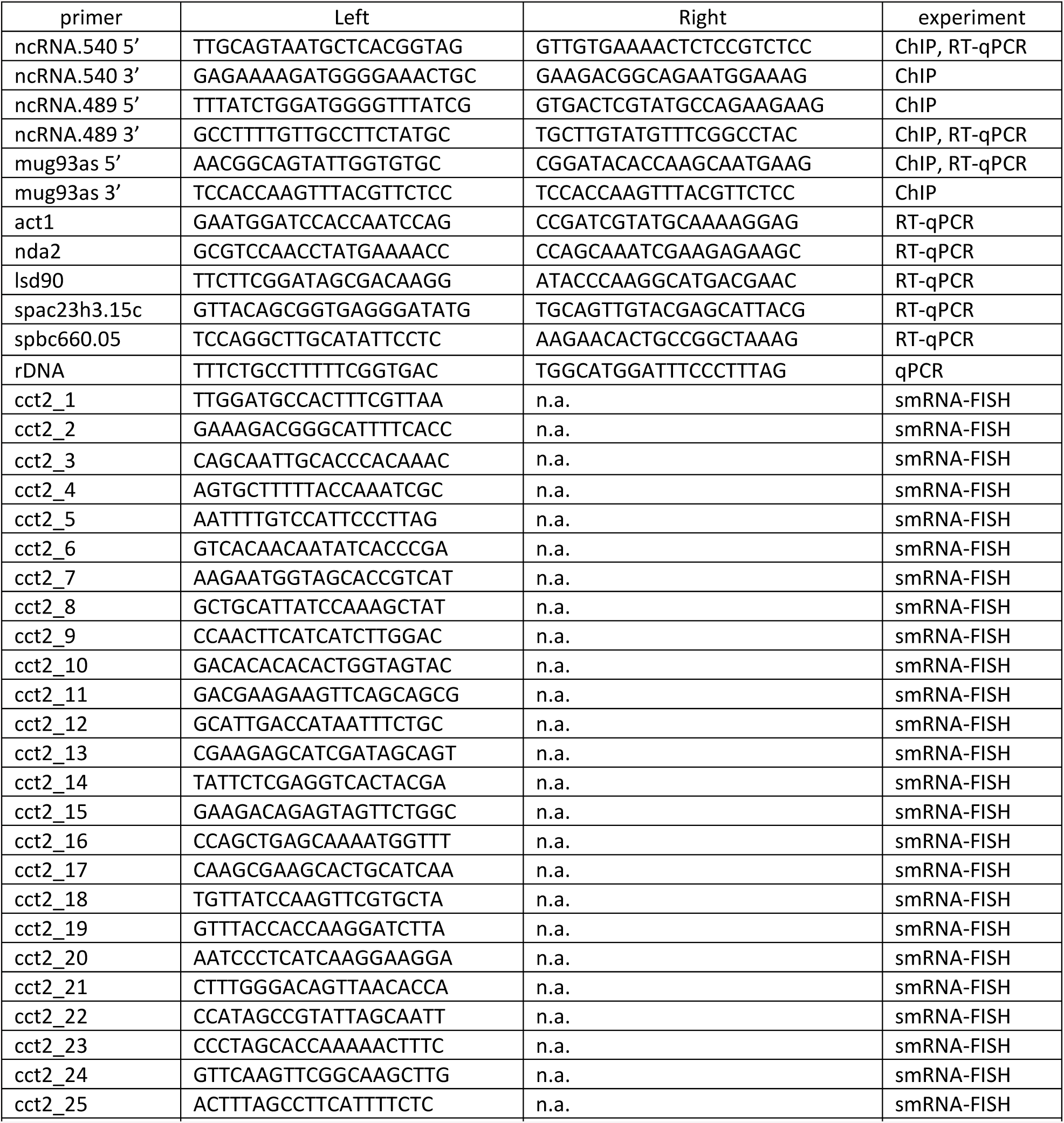
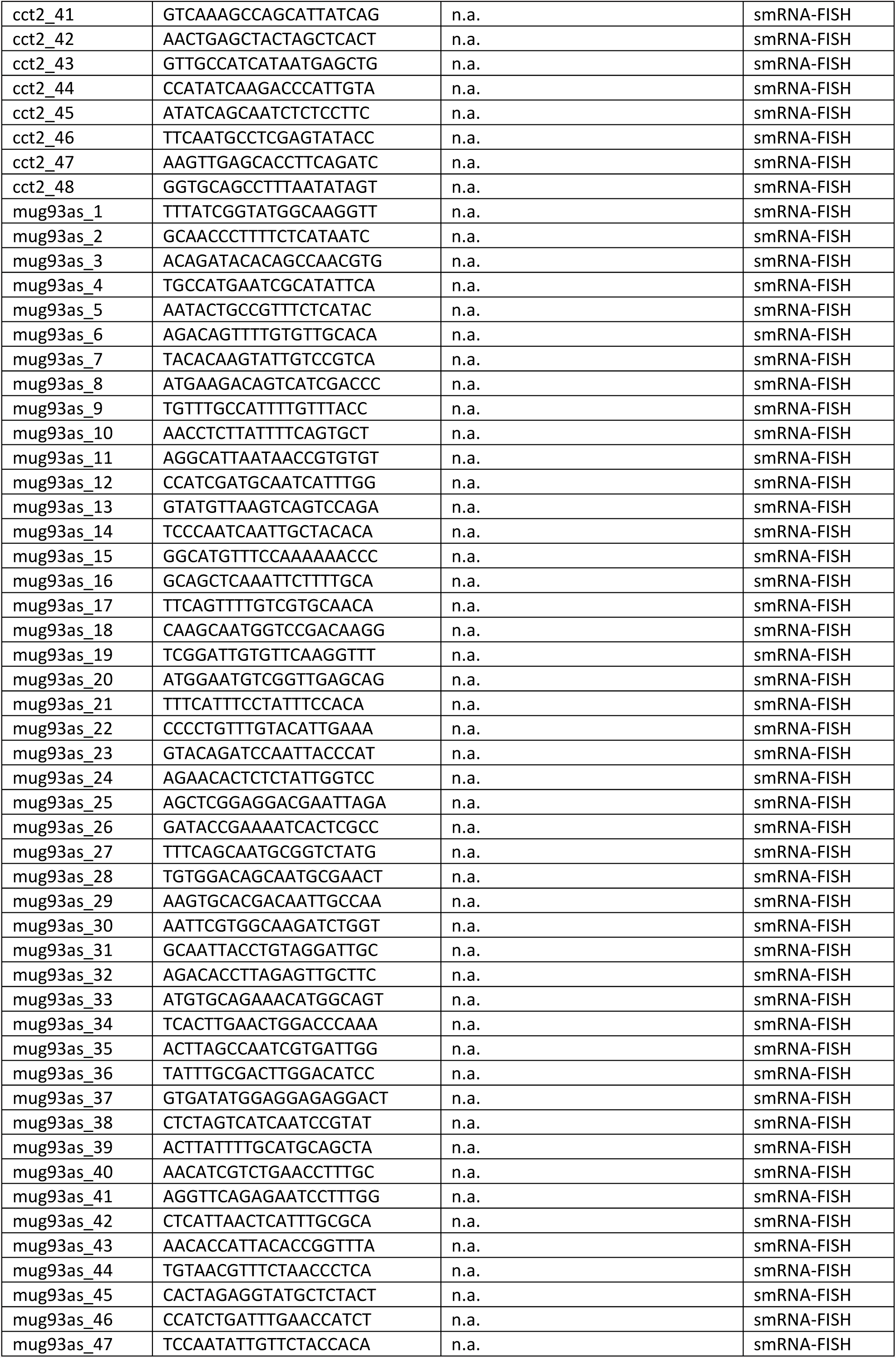
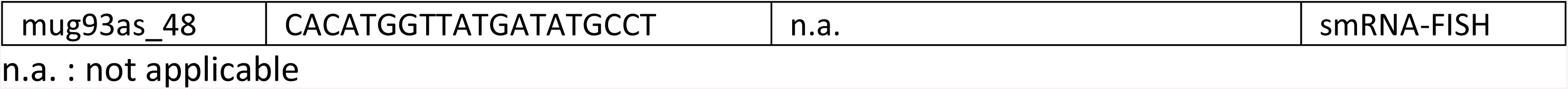
Primers and RNA-FISH probes used in this study

**Table S4.** Antibodies used in this study

| Application | Antibody |
| --- | --- |
| IF -α tubulin | mouse monoclonal Tat1, Keith Gull |
| IF Rrp6-MYC | mouse monoclonal #9E10 Thermo Fisher |
| IF Dis3-HA | mouse monoclonal #12CA5 Sigma-Aldrich |
| IF Gar2-GFP | rabbit polyclonal #A11122, Life Technologies |
| ChIP RNA Pol II (ser2P) | rabbit polyclonal #ab5095 ChIP-grade, abcam |
| ChIP anti-HA | Monoclonal 16B12 (anti-HA.11, Covance) |
| ChIP anti-PK | Monoclonal anti-PK (V5) Abd Serotec MCA1360 |
| Chromosome spreads anti-Brn1-HA<sub>6</sub> | Monoclonal 16B12 (anti-HA.11, Covance) |

**Table S5.** Microarray analysis G1-arrested cells

| id | tx_ID | seqName | strand | first | last | type | name | mappedType | source | Brn1_repea<br>t01 | Brn1_rep<br>eat02 | Brn1_rep<br>eat03 | Brn1(TEV)_<br>repeat01 | Brn1(TEV)_<br>repeat02 | Brn1(TEV)_<br>repeat03 | average<br>Brn1 | average<br>Brn1(TEV) | logRatio |
| --- | --- | --- | --- | --- | --- | --- | --- | --- | --- | --- | --- | --- | --- | --- | --- | --- | --- | --- |
| 5902 | ST5902 | 12 | - | 248597 | 249757 | ORF-T | YLR054C | bothEndsMapped | Manual | -3.48117 | -2.59869 | -2.09485 | -5.08599 | -5.03483 | -5.04824 | -2.34677 | -5.04154 | -2.69477 |
| 4501 | ST4501 | 5 | - | 52349 | 52485 | CUTs | CUT544 | mapped3 | Automatic | -4.43256 | -3.21128 | -4.46924 | -5.30533 | -7.65787 | -5.12852 | -3.84026 | -6.39319 | -2.55293 |
| 151 | ST0151 | 2 | + | 407335 | 410687 | ORF-T | YBR083W | bothEndsMapped | Manual | 0.16666 | 0.53404 | -4.84896 | -0.51932 | -4.52882 | -4.72394 | -2.15746 | -4.62638 | -2.46892 |
| 2138 | ST2138 | 12 | + | 105401 | 105545 | CUTs | CUT247 | mapped3 | Automatic | -5.83515 | -5.87913 | -3.45310 | -6.19907 | -8.21444 | -4.83796 | -4.66612 | -6.52620 | -1.86008 |
| 6627 | ST6627 | 14 | - | 777520 | 779200 | ORF-T | YNR074C | bothEndsMapped | Manual | -3.08368 | -2.43884 | -1.02514 | -1.83930 | -2.45676 | -4.04674 | -1.73199 | -3.25175 | -1.51976 |
| 3450 | ST3450 | 16 | + | 404243 | 404419 | SUTs | SUT407 | bothEndsMapped | Manual | -5.54183 | -3.48487 | -3.54037 | -3.15808 | -5.31097 | -4.52692 | -3.51262 | -4.91895 | -1.40633 |
| 5202 | ST5202 | 8 | - | 386606 | 386734 | CUTs | CUT651 | mapped3 | Automatic | -5.29801 | -4.13431 | -4.71461 | -6.09072 | -5.04489 | -6.31474 | -4.42446 | -5.67981 | -1.25535 |
| 5197 | ST5197 | 8 | - | 378126 | 379886 | ORF-T | YHR139C | bothEndsMapped | Manual | -5.14519 | -3.55026 | -3.84973 | -4.74112 | -4.43079 | -5.39266 | -3.70000 | -4.91173 | -1.21173 |
| 2492 | ST2492 | 13 | + | 239025 | 239225 | CUTs | CUT290 | mapped3 | Automatic | -5.31997 | -3.11358 | -5.14433 | -6.00678 | -5.04016 | -5.48125 | -4.12896 | -5.26070 | -1.13175 |
| 60 | ST0060 | 2 | + | 88041 | 88393 | other | snR56 | bothEndsMapped | Manual | -4.47023 | -3.18247 | -4.27987 | -4.60533 | -4.54478 | -5.16834 | -3.73117 | -4.85656 | -1.12539 |
| 3110 | ST3110 | 15 | + | 408138 | 408218 | SUTs | SUT363 | bothEndsMapped | Manual | -3.46131 | -4.57276 | -3.95585 | -5.75668 | -5.93836 | -4.82969 | -4.26431 | -5.38403 | -1.11972 |
| 6221 | ST6221 | 13 | - | 302549 | 302669 | CUTs | CUT765 | mapped5 | Automatic | -7.02135 | -3.68532 | -4.98378 | -5.93715 | -5.60098 | -5.29123 | -4.33455 | -5.44610 | -1.11155 |
| 5536 | ST5536 | 10 | - | 471068 | 471820 | ORF-T | YJR025C | bothEndsMapped | Manual | -1.24334 | -0.35086 | -0.47191 | -2.76667 | -1.39686 | -1.60625 | -0.41139 | -1.50156 | -1.09017 |
| 3983 | ST3983 | 3 | - | 178254 | 178478 | CUTs | CUT466 | mapped5 | Automatic | -3.98834 | -5.01892 | -5.04601 | -6.06312 | -6.05792 | -6.15247 | -5.03246 | -6.10519 | -1.07273 |
| 579 | ST0579 | 4 | + | 791956 | 792108 | CUTs | CUT070 | bothEndsMapped | Automatic | -6.16314 | -4.48993 | -3.56240 | -5.02902 | -4.95319 | -5.15140 | -4.02617 | -5.05229 | -1.02613 |
| 3123 | ST3123 | 15 | + | 435778 | 437386 | CUTs | CUT376 | mapped5 | Automatic | -5.25013 | -5.27972 | -4.93013 | -5.01997 | -4.09493 | -4.04544 | -5.10492 | -4.07018 | 1.03474 |
| 1643 | ST1643 | 9 | + | 281153 | 281257 | SUTs | SUT185 | bothEndsMapped | Manual | -4.88930 | -5.13821 | -4.99639 | -4.53378 | -4.60989 | -3.45173 | -5.06730 | -4.03081 | 1.03649 |
| 5190 | ST5190 | 8 | - | 364917 | 365365 | other | YHR130C, YHR131C | bothEndsMapped | Manual | -3.86027 | -3.20706 | -2.65851 | -4.53919 | -1.97958 | -1.57231 | -2.93278 | -1.77595 | 1.15684 |
| 3248 | ST3248 | 15 | + | 832339 | 832579 | other | snR8 | bothEndsMapped | Manual | -0.47182 | -2.33819 | 0.37779 | -1.43998 | 0.25944 | 0.27726 | -0.98020 | 0.26835 | 1.24855 |
| 6839 | ST6839 | 15 | - | 680566 | 680998 | other | snR36 | bothEndsMapped | Manual | -5.59490 | -5.29085 | -2.63283 | -5.40163 | -2.54971 | -2.79757 | -3.96184 | -2.67364 | 1.28820 |
| 1954 | ST1954 | 11 | + | 158572 | 159388 | ORF-T | YKL156W | bothEndsMapped | Manual | -2.25683 | -2.85398 | 0.25548 | -3.69957 | 0.27613 | -0.14455 | -1.29925 | 0.06579 | 1.36504 |
| 2658 | ST2658 | 13 | + | 731449 | 731553 | SUTs | SUT313 | bothEndsMapped | Manual | -5.65329 | -5.83973 | -6.45297 | -6.03445 | -4.64537 | -4.79716 | -6.14635 | -4.72127 | 1.42508 |
| 761 | ST0761 | 4 | + | 1401693 | 1402677 | ORF-T | YDR471W | bothEndsMapped | Manual | -0.84753 | -3.47101 | 0.70359 | -3.82080 | 0.60799 | 0.61579 | -1.38371 | 0.61189 | 1.99560 |
| 3679 | ST3679 | 1 | - | 199979 | 200419 | CUTs | CUT442 | bothEndsMapped | Automatic | -8.81809 | -3.38971 | -6.80481 | -6.61112 | -4.18029 | -1.69331 | -5.09726 | -2.93680 | 2.16046 |
| 6293 | ST6293 | 13 | - | 550109 | 551245 | ORF-T | YMR142C | bothEndsMapped | Manual | -4.31482 | -3.79717 | -3.54892 | -1.04337 | 0.75528 | -3.64919 | -3.67304 | -1.44695 | 2.22609 |
| 1911 | ST1911 | 11 | + | 15881 | 16201 | CUTs | CUT224 | mapped3 | Automatic | -2.37283 | -8.27058 | -7.37420 | -5.44901 | -6.21772 | -3.00349 | -7.82239 | -4.61060 | 3.21179 |

**Table S5. Microarrays analysis nocodazole-arrested cells**
| id | tx_ID | seqName | strand | first | last | type | name | mappedType | source | Brn1_repea<br>t01 | Brn1_repea<br>t02 | Brn1(TEV)_re<br>peat01 | Brn1(TEV)_rep<br>eat02 | average Brn1 | average<br>Brn1(TEV) | logRatio |
| --- | --- | --- | --- | --- | --- | --- | --- | --- | --- | --- | --- | --- | --- | --- | --- | --- |
| 4413 | ST4413 | 4 | - | 1319401 | 1319889 | ORF-T | YDR424C | bothEndsMapped | Manual | -0.23110 | -0.21460 | -3.16806 | -0.01457 | -0.22285 | -1.59131 | -1.36846 |
| 5671 | ST5671 | 11 | - | 179104 | 179464 | SUTs | SUT662 | bothEndsMapped | Manual | -0.48974 | -3.06247 | -2.87663 | -2.89802 | -1.77610 | -2.88732 | -1.11122 |
| 2088 | ST2088 | 11 | + | 554500 | 556036 | ORF-T | YKR059W | bothEndsMapped | Manual | -0.44989 | -0.05571 | -0.42022 | -2.19590 | -0.25280 | -1.30806 | -1.05527 |
| 2970 | ST2970 | 14 | + | 779124 | 781267 | ORF-T | YNR075W | bothEndsMapped | Manual | -1.97764 | -3.21846 | -1.58373 | -1.51300 | -2.59805 | -1.54836 | 1.04969 |
| 4957 | ST4957 | 7 | - | 735287 | 736026 | other | YGR122C-A | bothEndsMapped | Manual | -2.77460 | -1.14738 | -0.55634 | -0.79321 | -1.96099 | -0.67477 | 1.28622 |
| 6753 | ST6753 | 15 | - | 407822 | 408126 | other | snR9 | bothEndsMapped | Manual | 0.95456 | -1.43922 | 0.89781 | 1.47876 | -0.24233 | 1.18829 | 1.43062 |

